# Paired feed-forward excitation with delayed inhibition allows high frequency computations across brain regions

**DOI:** 10.1101/2021.03.13.435272

**Authors:** Alexandra S. Cao, Stephen D. Van Hooser

**Affiliations:** Department of Biology, Brandeis University, Waltham, MA 02454 USA; Volen Center for Complex Systems, Brandeis University, Waltham, MA 02454 USA; Sloan-Swartz Center for Theoretical Neurobiology, Brandeis University, Waltham, MA 02454 USA

**Keywords:** feed-forward, timing, cross-network computing, striate cortex, area 17, LGN

## Abstract

The transmission of high-frequency temporal information across brain regions is critical to perception, but the mechanisms underlying such transmission remain unclear. Long-range projection patterns across brain areas are often comprised of paired feedforward excitation followed closely by delayed inhibition, including the thalamic triad synapse, thalamic projections to cortex, and projections within hippocampus. Previous studies have shown that these joint projections produce a shortened period of depolarization, sharpening the timing window over which the postsynaptic neuron can fire. Here we show that these projections can facilitate the transmission of high-frequency computations even at frequencies that are highly filtered by neuronal membranes. This temporal facilitation occurred over a range of synaptic parameter values, including variations in synaptic strength, synaptic time constants, short-term synaptic depression, and the delay between excitation and inhibition. Further, these projections can coordinate computations across multiple network levels, even amid ongoing local activity. We suggest that paired feedforward excitation and inhibition provides a hybrid signal – carrying both a value and a clock-like trigger – to allow circuits to be responsive to input whenever it arrives.

## Introduction

In digital electronics, computations are synchronized by the presence of a digital clock trigger signal that indicates when each digital component should examine its inputs and perform a computation. The digital clock offers several advantages for digital circuits. First, it allows digital circuits to have short integration times, because the components do not need to consider information that has arrived at its inputs long in the past. Second, this short integration time allows the digital circuit to operate at high temporal speeds (many computations per second). Third, the digital clock allows synchronization of computations across different layers of digital components, because the arrival of related inputs is coordinated by the digital clock.

In neuronal circuits, several properties would seem to make coordinated and precise high-speed computation across layers impossible: the apparent lack of an equivalent clock, the electrical filtering that is performed by the cell’s membrane, and relatively slow synaptic time constants (von Neumann, 1963). However, we know that neural circuits do operate very quickly, as humans can make decisions within 200 milliseconds of the arrival of a visual stimulus (Thorpe et al., 1996; Sherwin et al., 2012). In the retina, ganglion cells can respond very well to temporal frequencies as high as 50 Hz (Frishman et al., 1987), and LGN neurons can reliably follow the high temporal frequencies of retinal ganglion cell inputs (Movshon et al., 2005), despite postsynaptic membrane filtering and synaptic depression. Further, in natural environments, stimuli may arrive at any time, at slow rates or fast rates, and circuitry must be ready to respond quickly regardless of when a previous stimulus arrived. Therefore, the brain must have mechanisms for quickly following high frequency inputs.

Many long-range connections across brain areas exhibit a peculiar motif: long-range feed-forward excitatory input is paired, after a short delay, with feed-forward inhibitory copy of the same input. At the retinogeniculate synapse, this inhibitory input is produced at a specialized dendro-dendritic synapse (the triad synapse) so that each feed-forward excitatory input is followed by a very fast and local inhibitory copy (Cox et al., 1998; Chen and Regehr, 2000; Blitz and Regehr, 2003, 2005). In other connections, such as those from thalamus to cortex, across regions of the hippocampus, or interareal connections, feed-forward excitatory inputs project to both principal neurons and inhibitory interneurons, and the delayed inhibitory input onto principal neurons arises from the interneurons that received feed-forward excitation (Buzsaki, 1984; Agmon and Connors, 1991; Swadlow and Gusev, 2000; Porter et al., 2001; Swadlow, 2003; Gabernet et al., 2005; Pouille et al., 2009; Yang et al., 2013; Rock and Apicella, 2015; Bhatia et al., 2019).

It has been previously recognized that the coordinated arrival of excitatory synaptic drive followed by inhibitory drive can result in very temporally precise postsynaptic action potentials and short integration windows (Pouille and Scanziani, 2001; Wehr and Zador, 2003; Blitz and Regehr, 2005; Mittmann et al., 2005; Higley and Contreras, 2006; Kremkow et al., 2010b; Cardin, 2018). Here in theoretical work we extend these findings to show that paired feedforward excitation and inhibition (FFEI) allows computation at very high temporal frequencies that “break” the membrane time constant barrier. Further, we show that this motif can synchronize computations across multiple hierarchical regions even when these regions have their own noisy ongoing local activity. Finally, we compare the limits of the temporal computations that are permitted in a low fan-in situation (1 paired synaptic input, such as at the triad synapse) and in a high fan-in situation (such as in input to cortex).

High temporal frequency transmission was possible over a wide range of synaptic parameters, including variations in synaptic strengths, synaptic time constants, synaptic depression, and delays between excitation and inhibition, suggesting that high temporal frequency transmission should be a characteristic of a variety of circuits that have this input structure. We conclude that paired feedforward excitatory and delayed inhibitory input is a hybrid signal, providing information about the value of the input while at the same time imposing a clock-like trigger signal that shortens temporal integration in a manner that allows high frequency computation and synchronization across networks or layers.

## Materials and Methods

Neural circuit simulations were performed in MATLAB (MathWorks, Natick, MA). Leaky integrate and fire neurons were modeled with the following differential equation, using the forward Euler method with a 0.1 ms time step (Lapicque, 1907; Abbott, 1999):

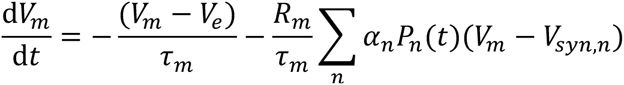

where *V_m_* is the membrane potential of the neuron, *V_e_* is the leak potential, *τ_m_* is the membrane time constant, *R_m_* is the membrane resistance, *P_n_*(*t*) is the synaptic conductance from synaptic input *n*, and *V_syn,n_* is the resting potential at the synapse for input *n*, and α*_n_* (= 1 for excitatory synapses) is a scaling coefficient for the synaptic current from input *n*. When *V_m_* reaches the threshold potential *V_thresh_*, the model neuron generates a spike and *V_m_* is set to the reset potential *V_reset_* (see Tables 1 and 2 for values).

**Table 1.**
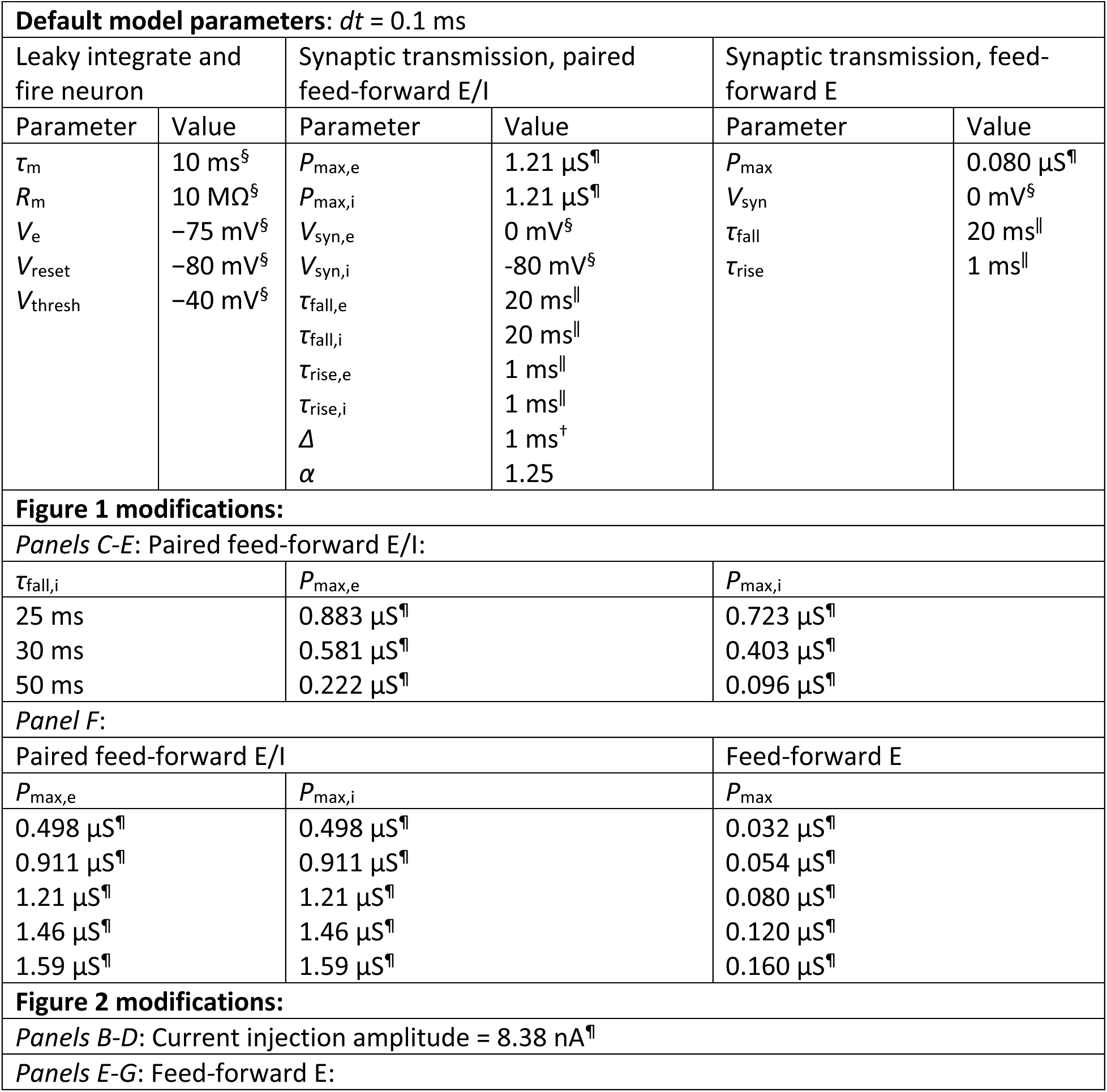
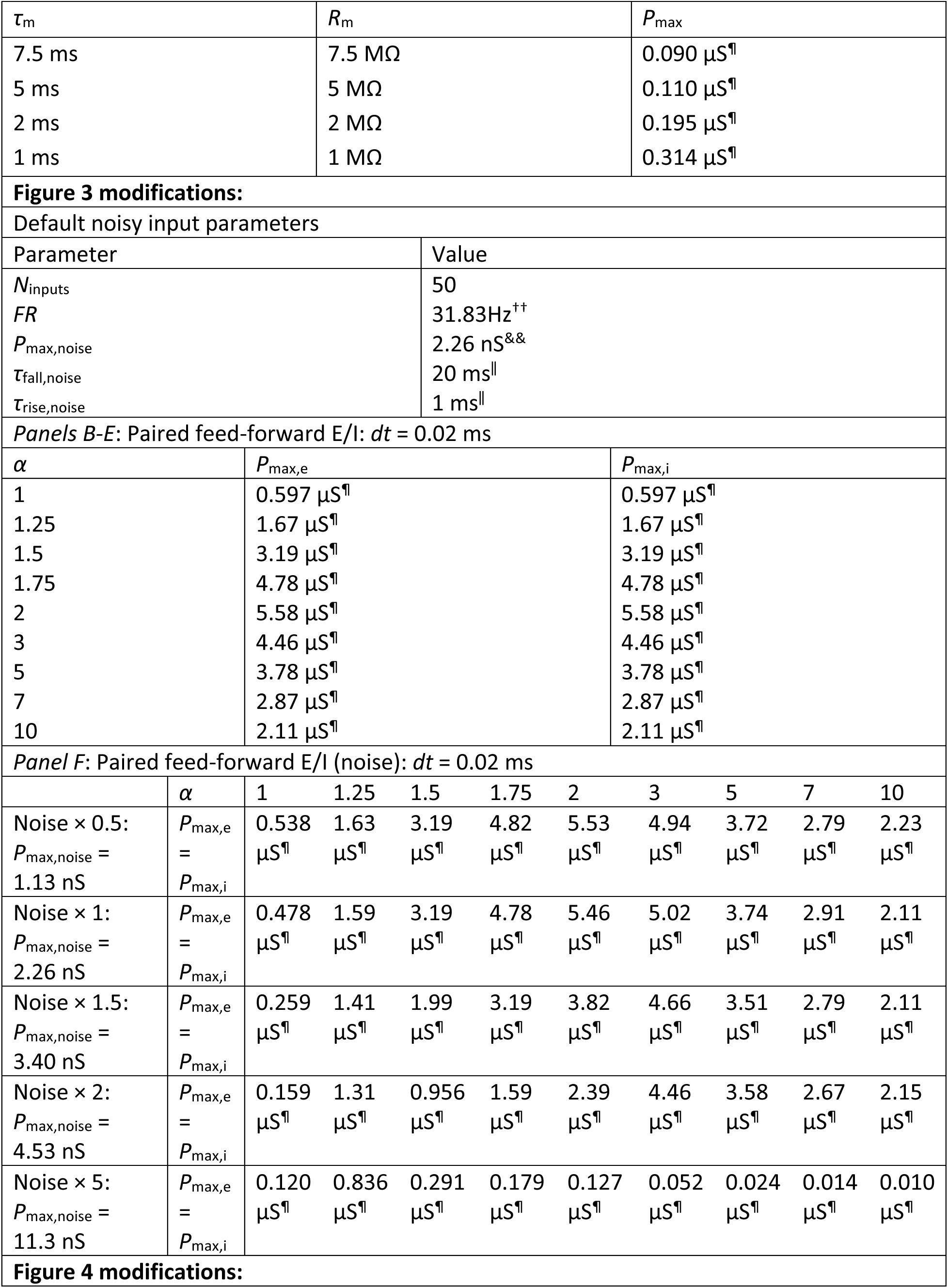
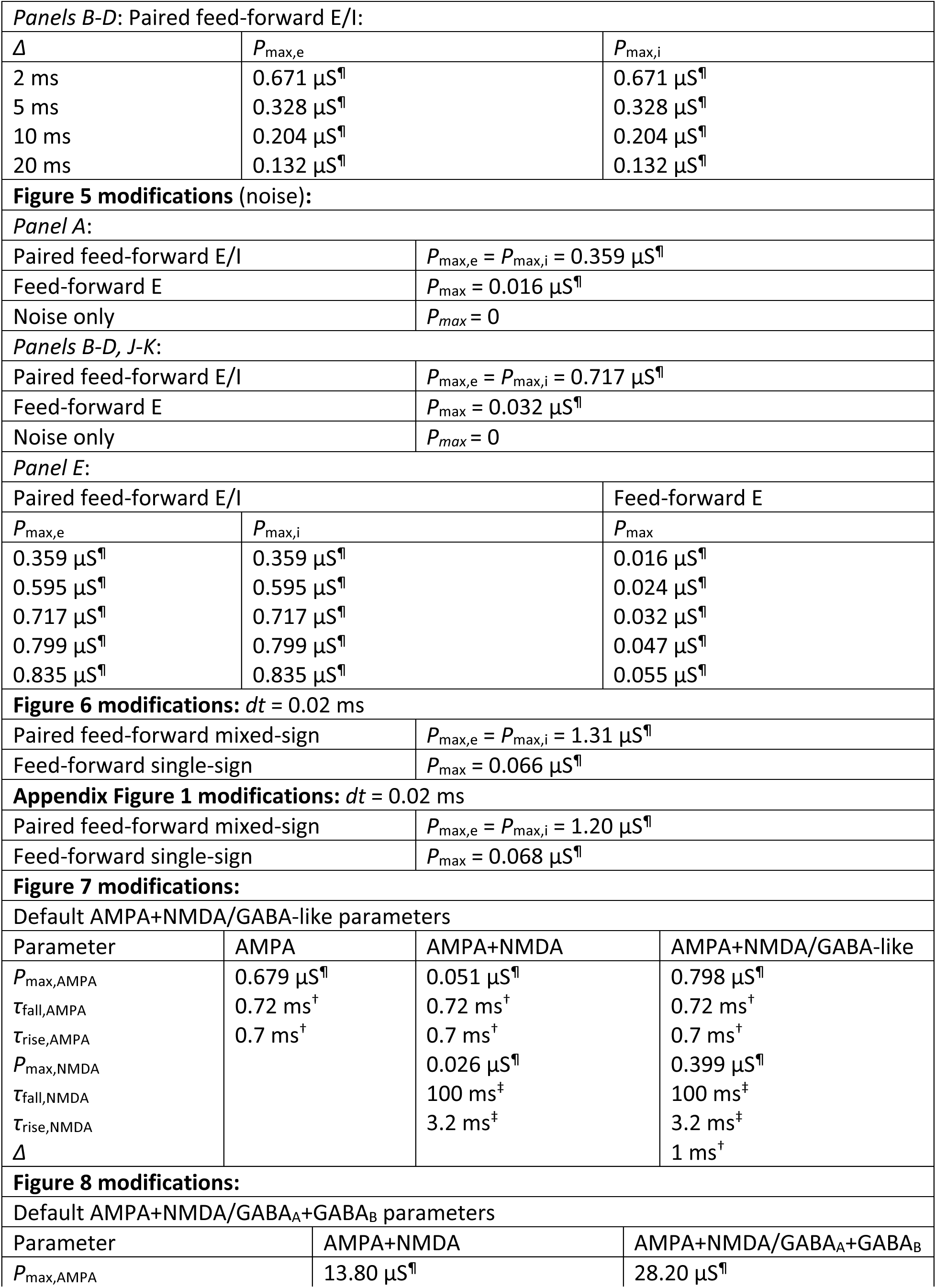
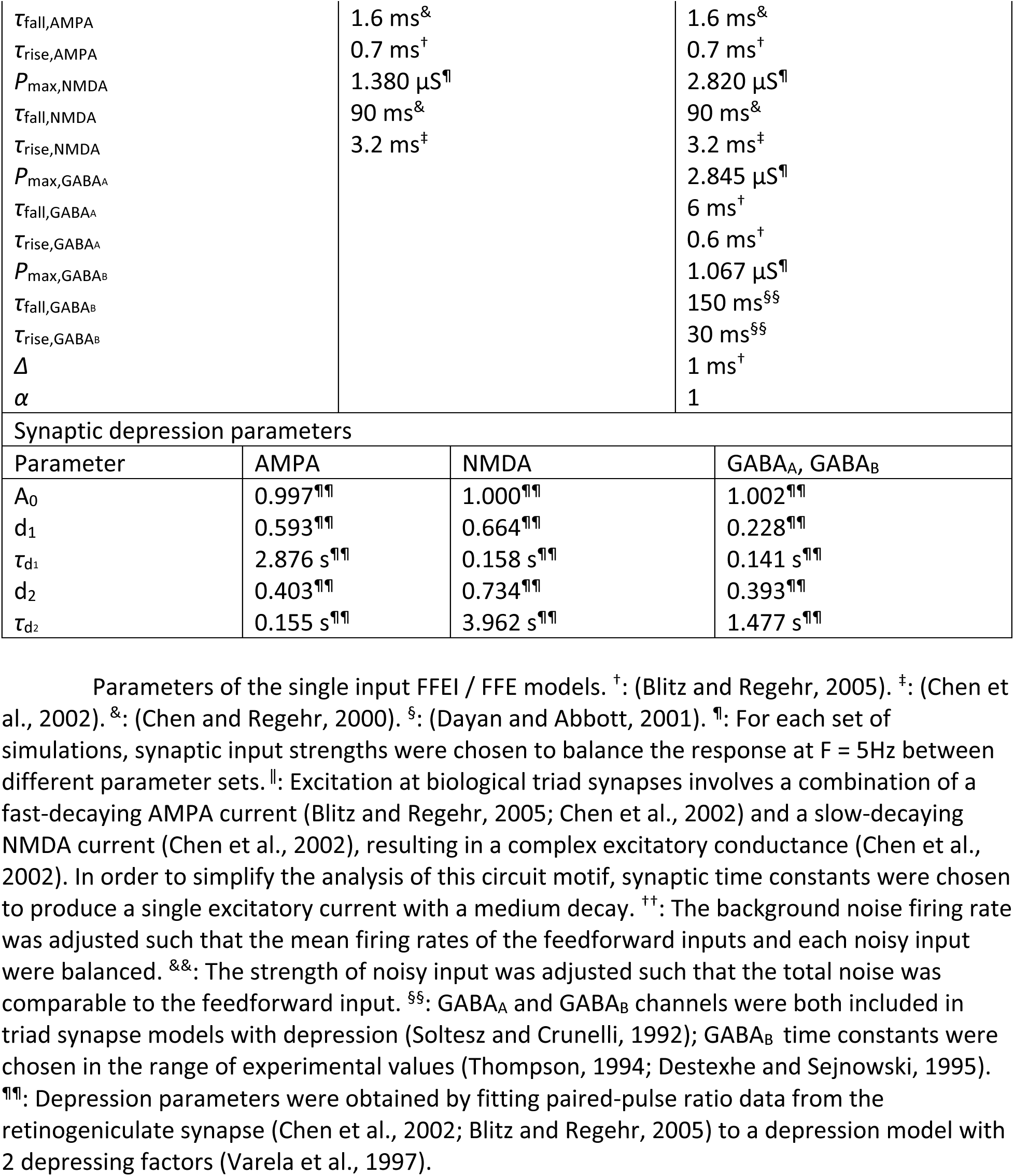
Parameters for triad synapse simulations.

**Table 2:** Parameters for LGN-cortex simulations.

| Default model parameters: $dt = 0.1$ ms | | | | |
| --- | --- | --- | --- | --- |
| Leaky integrate and fire neurons |  |  |  |  |
| Parameter | Interneuron (I) |  |  | Cortical neuron (E) |
| $\tau_m$ | 10 ms<sup>§</sup> | | | 10 ms<sup>§</sup> |
| $R_m$ | 10 MΩ<sup>§</sup> | | | 10 MΩ<sup>§</sup> |
| $V_e$ | -75 mV<sup>§</sup> | | | -75 mV<sup>§</sup> |
| $V_{reset}$ | -80 mV<sup>§</sup> | | | -80 mV<sup>§</sup> |
| $V_{thresh}$ | -48 mV<sup>◊</sup> | | | -40 mV<sup>§</sup> |
| Synaptic transmission |  |  |  |  |
|  | Paired feed-forward E/I |  |  | Feed-forward E |
| Parameter | LGN input to (E) | LGN input to (I) | (I) to (E) | LGN input to (E) |
| $P_{max}$ | 2 nS | 0.6 nS | 100 nS | 2 nS |
| $V_{syn}$ | 0 mV<sup>§</sup> | 0 mV<sup>§</sup> | -80 mV<sup>§</sup> | 0 mV<sup>§</sup> |
| $\tau_{fall}$ | 20 ms<sup>#</sup> | 20 ms<sup>#</sup> | 20 ms<sup>#</sup> | 20 ms<sup>#</sup> |
| $\tau_{rise}$ | 1 ms<sup>#</sup> | 1 ms<sup>#</sup> | 1 ms<sup>#</sup> | 1 ms<sup>#</sup> |
| $N_{inputs}$ | 100 | 100 | 1 | 100 |
| $\alpha$ | | | 1.25 | |
| Figure 9 modifications: |  |  |  |  |
| *Panels C-E:* (I): $\tau_m = [5$ ms, 6 ms, 10 ms, 20 ms, 50 ms, 100 ms] | | | | |

Synapses were modeled as a difference of exponentials, with two time constants (Dayan and Abbott, 2001):

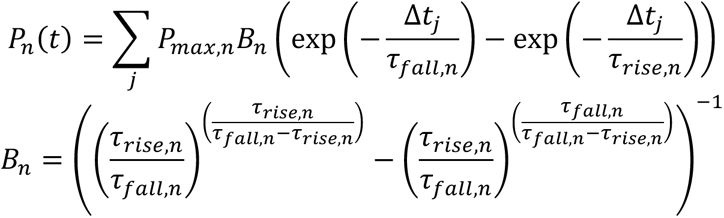

where *P_max,n_* is the maximum conductance of the synapse, *B_n_* is a normalization factor to ensure that the peak value of *P_n_*(*t*) is *P_max,n_*, Δ*t_j_* is the time between *t* and the *j*th spike of synaptic input *n*, *τ_fall,n_* is the fall time constant, and *τ_rise,n_* is the rise time constant. (see Tables 1 and 2 for values).

### Simulating inputs to neural circuits

Temporal information was generated using a Poisson process with a rectified sinusoidal firing rate:

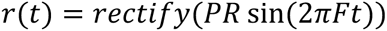

where *PR* is the peak firing rate set to 100 Hz for all simulations, *F* is the input modulation frequency, and rectify(x) = 0 if x < 0, and rectify(x) = x for x≥0. For each set of simulations, temporal information was presented to our simulated neural circuits for a fixed length of time, independent of *F*.

In some of our simulations, we provided random background activity as an additional excitatory synaptic input. This activity was modeled as *N_inputs_* noisy Poisson inputs with a uniform firing rate *FR* (see Table 1 for values). *FR* was calculated such that, for each noisy input, the probability of seeing 1 background spike during an input cycle is equal to the probability of a feedforward input spike over one cycle (that is, the mean firing rate of the feedforward inputs and each noisy input were balanced):

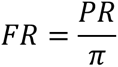

### Calculating the power of spiking responses at a particular frequency

Spiking responses were converted to a firing rate per time step (Hz), and the following equation was used to calculate the Fourier coefficient magnitude at a particular frequency (Crawford, 1968):

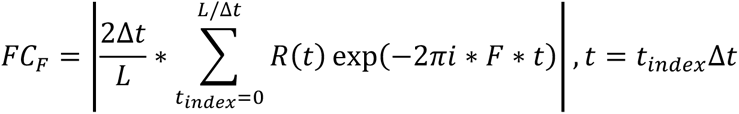

where Δ*t* is the time step size in seconds, *L* is the length of the spiking response in seconds, *t_index_* is time as an integer index, *R*(*t*) is the spiking response in Hz, and *F* is a particular frequency.

To normalize the power of spiking responses at a particular frequency, we used the average power calculated over all frequencies (*FC_avg_*) as a normalization factor. For a time interval of length *L* in time steps of Δ*t*, 1/Δ*t* and 1/*L* are the maximum and minimum frequencies that can be represented, respectively. Therefore, *FC_avg_* was determined by averaging *FC_F_* over *F* = 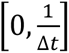 in intervals of 1/*L*. The normalized frequency response was then expressed as *FC_F_*/*FC_avg_*. In cases where *FC_avg_* = 0, the normalized frequency response was set to 0.

Unless otherwise stated, *FC_F_*, *FC_avg_*, and *FC_F_*/*FC_avg_* were each averaged over 10 trials for each model.

### Balancing excitation and inhibition in triad synapse models

When varying the inhibitory fall time constant (*τ_fall,i_*) in our triad synapse models, we scaled *P_max,i_* relative to *P_max,e_*, such that the total excitation and inhibition are balanced:

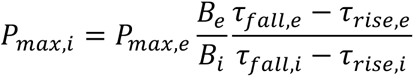

The above equation was derived by first integrating *P_e_*(*t*) and *P_i_*(*t*) for a single spike at *t* = 0 over the interval *t* = [0, ∞], as a measure of the total conductance due to a single spike. *P_max,i_* values were then obtained by setting the total excitation equal to the total inhibition, and then solving for *P_max,i_* given a set of time constants and excitatory parameters.

### Determining the expected output of an exclusive-or computation

In Figure 6, we examine the output of a circuit designed to perform an exclusive-or (XOR) computation of 2 inputs. The expected output of the circuit was determined by performing the XOR computation on sliding windows of length *t_bin_* = 10 ms or 5 ms. If only one of the inputs fires during the time interval 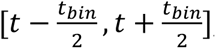, then output is expected at time *t*.

### Simulating depression at the retinogeniculate synapse

Paired-pulse ratio data for AMPA and NMDA (Chen et al., 2002) and for GABA (Blitz and Regehr, 2005) in the retinogeniculate synapse were fitted to the following depression model (Varela et al., 1997):

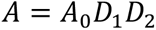

In the above equation, *A* is a scaling factor on the amplitude of the synaptic current generated by each presynaptic spike, *A*_0_ is the undepressed scaling factor, and both *D*_1_ and D_2_ are depressing factors with default values of 1. Following each presynaptic spike, *D_i_* is set to *D_i_d_i_*, and *D_i_* decays back to 1 with a time constant 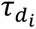 (see Table 1 for values and Fig. 8 supp. 1 for fits).

The code for the paper is at https://github.com/VH-Lab/vhlab-ffei-matlab (Note to reviewers and editors: this is currently private but will be made public on acceptance)

### Key Resources Table

| Reagent type (species) or resource | Designation | Source or reference | Identifiers | Additional information |
| --- | --- | --- | --- | --- |
| Software, algorithm | Matlab | The MathWorks, Natick, MA | RRID:SCR_001622 |  |
| Software, algorithm | GitHub | GitHub | RRID:SCR_002630 |  |

### Sex as a variable

Not applicable to the computational models studied here.

## Results

Our goal was to understand if there was any potential advantage of the paired feedforward excitatory and inhibitory (FFEI) connections that are commonly observed in projections across brain regions. We analyzed the problem in two regimes. First, we analyzed a low convergence situation (N_inputs_ = ∼1), inspired by the triad retinogeniculate synapse in the LGN. Here, we discovered that FFEI projections allowed feed-forward computations at much higher frequencies than if feedforward excitatory (FFE) projections were employed alone. In order to focus specifically on the difference between FFEI projections and FFE projections, we first explored models that were inspired by the LGN triad synapse but were much simpler, to demonstrate the phenomena and how they depend on the inhibitory delay and properties of the inhibitory synapse. Next, we explored a more realistic model of the triad synapse that includes synaptic depression of excitatory and inhibitory synapses, and showed that the principles of the very simple models apply under these conditions. Finally, we examined a high convergence situation (N_inputs_ = ∼50-150) such as is found at thalamocortical and intercortical projections.

### Paired feedforward excitatory and inhibitory inputs allow transmission of temporal information at high frequencies

We began with a simple model inspired by the retina-LGN triad synapse that is particularly common at synapses between retinal X cells and LGN X cells (Koch, 1985; Hamos et al., 1987; Lam et al., 2005; Bickford, 2019). At these synapses, excitatory retinal input is provided to an LGN neuron and to an adjacent terminal bouton of the dendrite of an interneuron. The interneuron dendrite, which is electotonically isolated from its soma (Morgan and Lichtman, 2020), also contacts the LGN cell at very nearly the same location as the excitatory cell, providing an only slightly-delayed inhibitory input to the LGN neuron (Cox et. al, 1998). To build our model, we included a feedforward glutamatergic excitatory synaptic current with temporal dynamics that were set to be between the fast-decaying AMPA current and the slower-decaying NMDA current observed at triad synapses (Chen et al., 2002). Fast GABAergic currents were included with a 1 ms delay, also following experimental measurements (Blitz and Regehr, 2003, 2005). While the total synaptic conductance was varied, the maximal conductances of the excitatory and inhibitory synapses were selected so that the total conductance (the area under the curve) of excitation and inhibition were identical; that is, the total integrated conductance of excitation and inhibition were balanced.

To assess the transmission of temporal information in this model, we simulated the spiking responses of a leaky integrate-and-fire neuron to input spikes generated via a Poisson process with a rectified sinusoidal firing rate (Fig. 1AB). The rate at which spikes were generated varied in time sinusoidally from 0 spikes/sec to 100 spikes/sec, and the frequency of the sinewave was termed the modulation frequency. At each 0.1ms, a random number (0-1) was drawn, and a spike was generated if the random number was less than the product of the rate and the bin size (0.1ms). While the FFE configuration produces extra spikes that are outside the sinusoidal input modulation, output spikes in the FFEI input configuration are well locked to the sinusoidal input modulation. We examined transmission of the FFEI projection over a range of inhibitory synapse fall time constants that varied from 20 to 50 ms, and compared its transmission to the case where only feedforward excitatory (FFE) input was provided. Because changing the fall time constant of the inhibitory synaptic current, while keeping integrated E and I conductance fixed, alters the fraction of the excitatory conductance that arrives earlier in the postsynaptic response, the overall responsiveness of the cell also varies as we adjusted the inhibitory synapse fall time constant. To compare equally responsive models, we adjusted the feedforward synaptic weight so that each model generated a response of about 75 Hz when driven with an input modulating frequency of 5 Hz.

**Figure 1.**
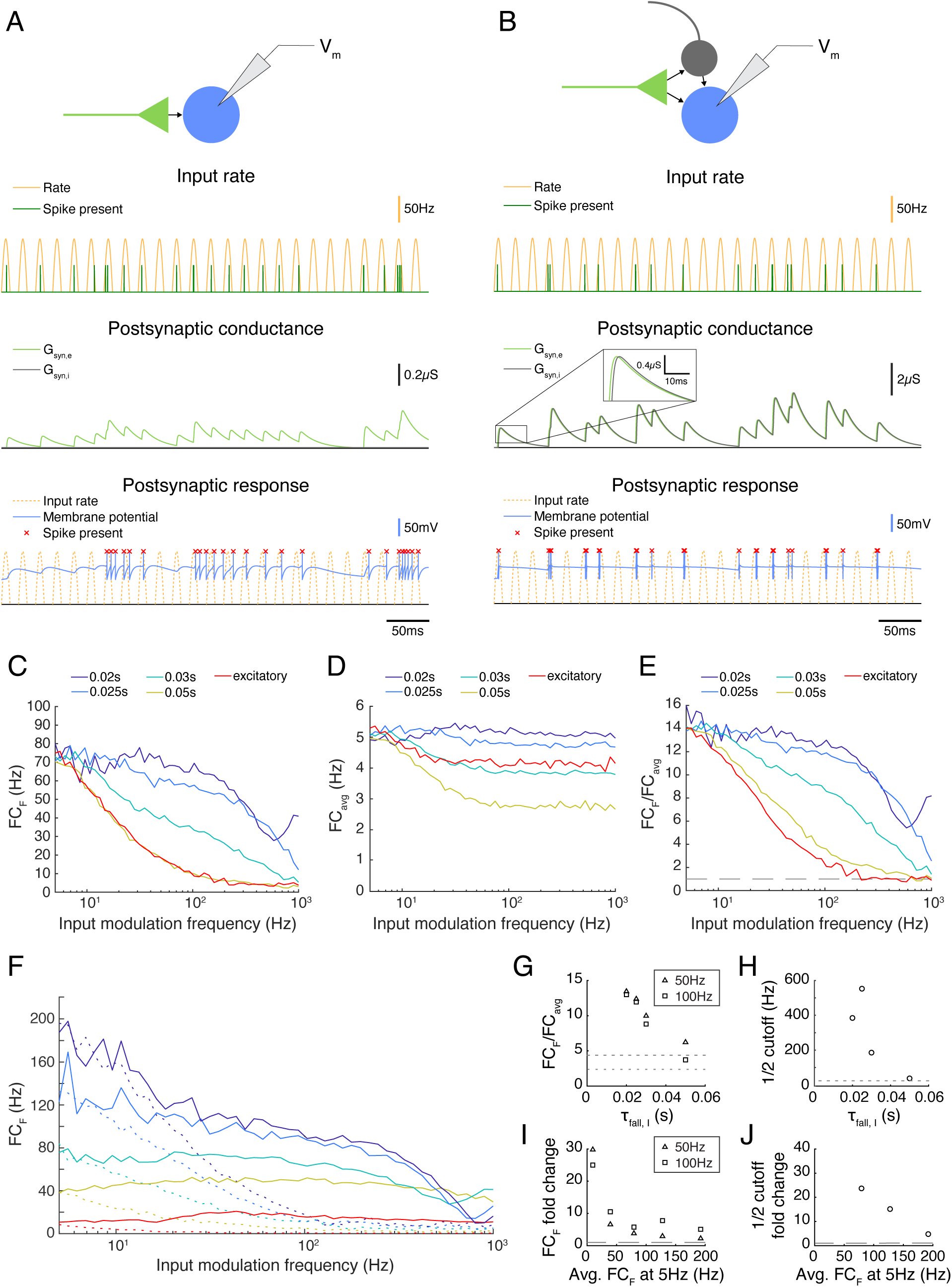
Paired feedforward excitation and delayed inhibition in a triad synapse model allows transmission of higher frequencies than feedforward excitation alone. **(A) Top**: A feedforward excitatory (E) model circuit with a single RG-like cell (light green) providing excitatory input to an LGN-like neuron (blue). **Bottom**: A 0.5 s simulation of the model circuit with a rectified sinusoidal input firing rate (yellow) with a peak rate (PR) of 100 Hz and an input modulation frequency (F) of 50 Hz; presynaptic spikes were generated with a Poisson process (dark green). The postsynaptic conductance (light green) and the membrane potential of the LGN cell (blue) are simulated in response to the feedforward input. Output firing times are marked (red x), and the postsynaptic response is compared to the underlying input characteristics (dashed yellow). **(B)** Same, but for a paired feedforward excitatory/inhibitory (E/I) model circuit with a single feedforward cell (light green) providing excitatory input to an output neuron (blue) and to an interneuron (gray), which in turn provides delayed inhibitory input (1 ms) to the output neuron. **(C)** For a range of input modulation frequencies F, the Fourier coefficient of the output at F (FC_F_) was averaged over 10 trials for triad synapse models with different inhibitory conductance characteristics. Legend entries indicate the length of the inhibitory time constant τ_fall,I_ for each paired feedforward E/I model; “excitatory” indicates the feedforward E model. Synaptic strengths were adjusted so that Fourier coefficients at input modulation 5 Hz were approximately 75 Hz. **(D)** Mean Fourier coefficient of the output over all frequencies, including those that are much higher and lower than the modulation frequency. **(E)** Fourier coefficient of the output at F divided by the mean overall power (FC_F_/FC_avg_). Dashed line indicates FC_F_/FC_avg_ = 1. **(F)** Same as C, except that synaptic strength P_max_ was adjusted to produce a range of Fourier coefficient values at an input modulation frequency at 5 Hz (weak to strong: red to dark blue). Solid lines indicate the feedforward E/I model, and dashed lines indicate the feedforward E model. In all cases, paired E/I input resulted in greater transmission at high input modulation frequencies than E input alone. **(G)** The influence of the inhibitory synaptic fall-off time constant τ_fall,I_ on transmission FC_F_/FC_avg_, measured at F = 50 Hz (triangles) and 100 Hz (squares). Dashed lines indicate feedforward E model at 50 Hz (top) and 100 Hz (bottom). The paired E/I configuration exhibits greater transmission over a range of inhibitory synaptic time constants. **(H)** The influence of the inhibitory synaptic fall-off time constant τ_fall,I_ on the input modulation frequency at which FC_F_/FC_avg_ is ½(FC_F_/FC_avg_ at F = 5 Hz). Dashed line indicates the ½ cutoff for the feedforward E model. **(I)** For each paired feedforward E/I model shown in (F), the FC_F_ fold change with respect to the feedforward E model of matching response at F = 5 Hz for F = 50 Hz (triangles) and F = 100 Hz (squares). Dashed line indicates a fold change of 1. **(J)** For each paired feedforward E/I model shown in (F), the ½ cutoff fold change with respect to the feedforward E model of matching output power (denoted by line style and FC_F_ at F = 5 Hz) was calculated as FFEI/FFE. For the lowest firing rates, ½ cutoff was greater than 1000 Hz in the FFEI case (out of range). Dashed line indicates a fold change of 1. In all, paired E/I input allowed transmission of higher input modulation frequencies over a wide range of inhibitory synaptic time constants and for a wide range of synaptic input strengths.

FFEI projections demonstrated a remarkable ability to transmit high frequency information. To measure transmission quality at each input frequency, we calculated the Fourier coefficient (FC) of the spiking response at the input modulation frequency (FC_F_, Fig. 1C). The Fourier coefficient is higher when spikes are well locked to the stimulus, and drops as the number of spikes fired per cycle becomes lower or becomes statistical (that is, with some missed cycles). Projections with only excitatory synapses were filtered strongly at input modulation frequencies ranging from 20 – 100 Hz. In contrast, FFEI projections with inhibitory time constants of 20 ms transmitted with nearly equal fidelity from 5 – 100 Hz, and did not reach a 50% reduction in Fourier coefficient responses until the input modulation frequency reached about 400 Hz.

While the Fourier coefficient at the input modulation frequency (FC_F_) measures the output transmission at that single frequency, it did not provide information about how much of the total output of the cell was concentrated exclusively at the input modulation frequency. For example, if the output cell increased its overall response only tonically, then FC_F_ would still exhibit an increase. To examine the fraction of the output cell’s total frequency response that was at the input modulation frequency, we divided FC_F_ by the average Fourier coefficient calculated over all frequencies, including frequencies much higher and lower than the input modulation frequency, (Fig. 1D) to produce a normalized frequency response measure (Fig. 1E). This analysis indicated that FFEI projections transmitted information that was highly specific to the input modulation frequency over a wide frequency range 5 – 100 Hz, while the specificity of information transmitted by excitatory-alone projections fell off rapidly with frequency.

The ability to provide specific output at the input modulation frequency was highly related to the inhibitory fall time constant. When the inhibitory fall time constant was short, then the advantage of the FFEI configuration was greater. When the time constant is small (20 ms), then the inhibition comes right on the heels of the excitation so that the excitation cannot drive the cell for a long time. When there is no inhibition, the excitation can drive the cell for a long time, which means less of the response is at the input modulation frequency. When there is a long time constant of inhibition (50 ms), then the inhibition is more spread out in time and the behavior is more intermediate, although more single feedforward input spikes are missed by the output neuron than in the short time constant case (**Appendix Figure 1**).

We plotted the normalized output responses for input modulation frequencies of 50 and 100 Hz in Fig. 1G. For an inhibitory fall time constant of 20 ms, the postsynaptic neuron produced output at the input modulation frequency that was more than 12 times the amount of average output at other frequencies. As the inhibitory fall time constant was increased, the output neuron behaved more like the excitatory-only configuration, as one would expect. The ½ cutoff frequency – that is, the input modulation frequency at which the normalized output dropped to ½ of the value it showed for a 5 Hz input – was also highly dependent on the inhibitory fall time constant (Fig. 1H).

It was possible that the relative advantage of FFEI transmission compared to FFE was not general, but instead was specific to the particular levels of synaptic input strengths that were chosen in order to drive output responses of about 75 Hz at a 5 Hz input modulation frequency. To examine this possibility, we compared FFEI and FFE models that were matched at 5 Hz input modulation but over a wide variety of output response levels (Fig. 1F). In all cases, the FFEI projection responses at the input modulation frequency were much stronger than those of the FFE projection over a wide range of input modulation frequencies. To quantify these changes, we examined the fold change of transmission of FFEI compared to FFE at input modulation frequencies of 50 or 100 Hz for these different levels of synaptic drive (Fig. 1I). For all of these levels of synaptic drive, the FFEI projection produced at least 2 times as much drive at 50 and 100 Hz compared to FFE alone. The ½ cutoff frequency was also more than 4 times higher for FFEI compared to FFE (Fig. 1J).

### Paired feedforward excitatory and inhibitory inputs allow effective transmission at frequencies that are highly filtered by the output cell’s membrane properties

The ability of the output cell to reliably follow very high input frequencies was surprising because the membrane properties of neurons act as a low-pass filter, and high frequency current fluctuations are not well transmitted to membrane voltage. We compared the ability of the output neuron to follow synaptic input provided through either FFEI or FFE projections with the ability of the output neuron to follow direct sinusoidal membrane current injections of a fixed amplitude (Fig. 2).

**Figure 2.**
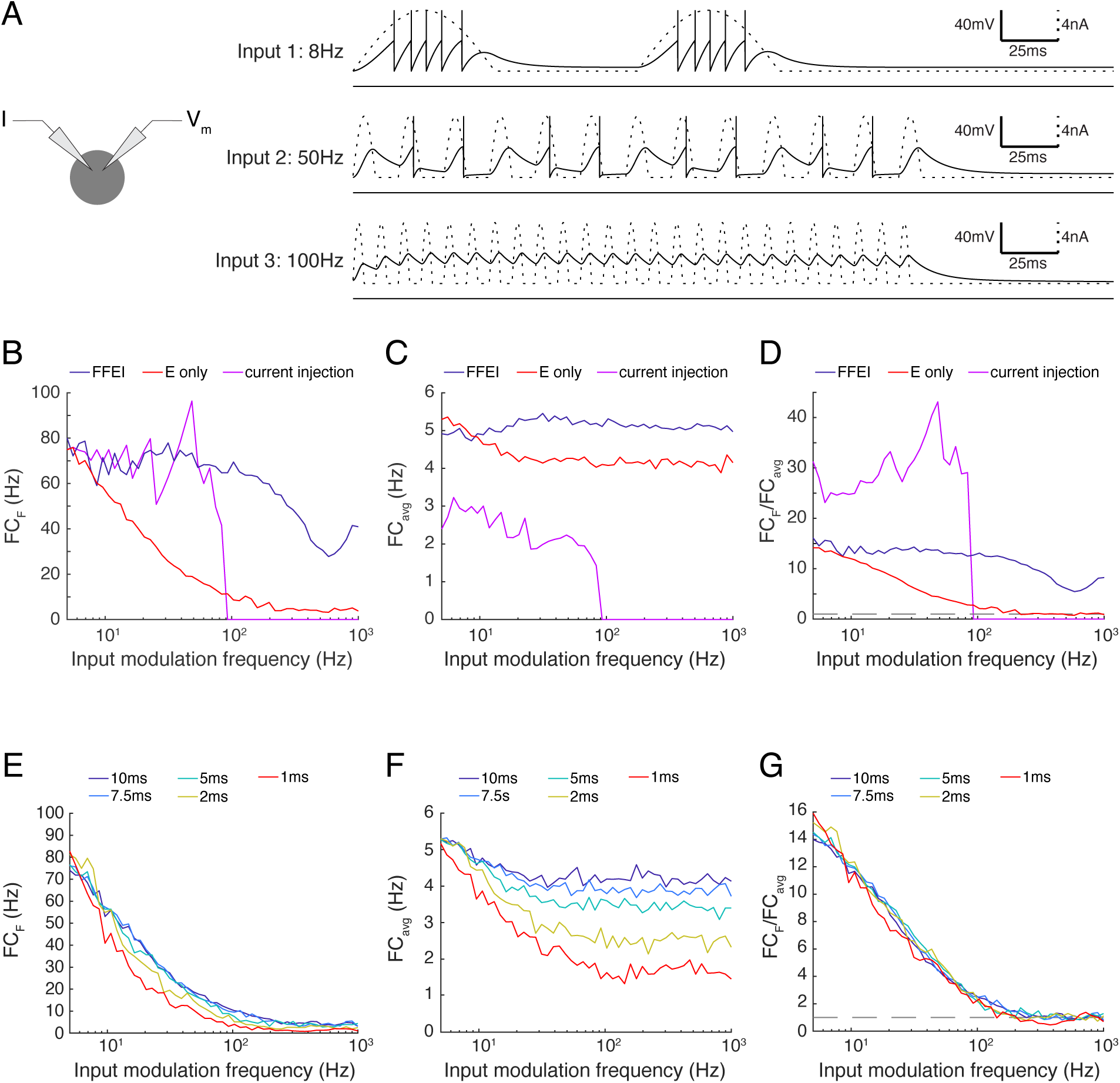
Paired feed-forward E/I inputs allows transmission of higher input modulation frequencies compared to direct sinusoidal current injection; enhanced high frequency transmission is not attributed to changes in the effective membrane time constant. **(A)** Left: A model cell receiving direct injections of rectified sinusoidal currents at various frequencies. As input modulation frequency increases, the ability of the cell to follow the input decreases, and the cell is totally unable to spike in response to the input modulation frequency of 100 Hz. **(B)** Fourier coefficient of spiking output for a range of input modulation frequencies F for direct current injections (magenta), paired feedforward E/I model with τ_fall,I_ = 20ms (blue), and for the feedforward E model (red). **(C)** Same, except mean responses over all Fourier frequencies is shown (FC_avg_). **(D)** The normalized output response (FC_F_/FC_avg_). Dashed line indicates FC_F_/FC_avg_ = 1. When firing rates are matched at input modulation frequency of 5Hz, direct current injections provide more selective transmission for lower frequencies but paired feedforward E/I connections allow the cell to follow signals at higher frequencies than direct current injection. **(E)** Adjustment of the membrane time constant over a 10-fold range did not allow high frequency transmission. Fourier coefficient of feedforward E model output for a range of input modulation frequencies F. Legend entries indicate the membrane time constant τ_m_ for each feedforward E model. **(F)** Mean feedforward E model responses over all Fourier frequencies. **(G)** The normalized output response (FC_F_/FC_avg_). Dashed line indicates FC_F_/FC_avg_ = 1. High frequency transmission in the feedforward E model is not substantially enhanced by decreased τ_m_.

As before, we calibrated the output of the neuron to hit a target rate (75 Hz) at a 5 Hz input modulation frequency. For the FFEI and FFE simulations, we did this by setting the synaptic weight, and for the current injection simulations, we did this by setting the amplitude of the sinusoidal current injection. We compared the raw Fourier coefficient measured at the input modulation frequency (Fig. 2B), the overall response across all frequencies (Fig. 2C) and the normalized frequency response to measure the specificity of the output response to the input modulation frequency (Fig 2D). As expected, the cell followed the sinusoidal current injection with substantial fidelity until a critical frequency was reached, after which the input fluctuations became too small to generate action potentials. This resulted in a rapid drop in the ability of the cell to follow the output frequencies. By contrast, FFEI inputs continued to drive responses at high frequencies, well beyond the point at which the sinusoidal current could drive spikes. While the critical frequency at which spiking stops for direct current injection does depend upon the amplitude of the current used, the take-home point here is that FFEI projections drive responses at higher input frequencies when responses to FFEI and direct current injections are matched at low frequencies.

The addition of an inhibitory conductance in the FFEI case will necessarily reduce the effective membrane time constant, and one may ask whether the increased transmission is primarily due to the reduced time constant. To explore this, we modified the membrane time constant of our model neuron over a 10-fold range, and examined the influence of the passive membrane time constant on the temporal frequency of transmission (Fig. 2EFG). We observed very small influences of modifying the membrane time constant, indicating that a passive adjustment of the membrane time constant cannot account for increased high frequency transmission observed here.

At first glance, it seems almost paradoxical that inputs delivered via FFEI projections can apparently “break the limit” of the membrane time constant. To gain intuition into the mechanism of action of FFEI projections, we examined synaptic currents generated by a 100 Hz train of input spikes when delivered either by FFE or FFEI projections (Fig. 3A). To study the influence of different amounts of inhibition on transmission, we also varied the relative weight (total area under conductance curve per presynaptic spike) of the inhibitory conductance to the excitatory conductance, and called this quantity α.

**Figure 3.**
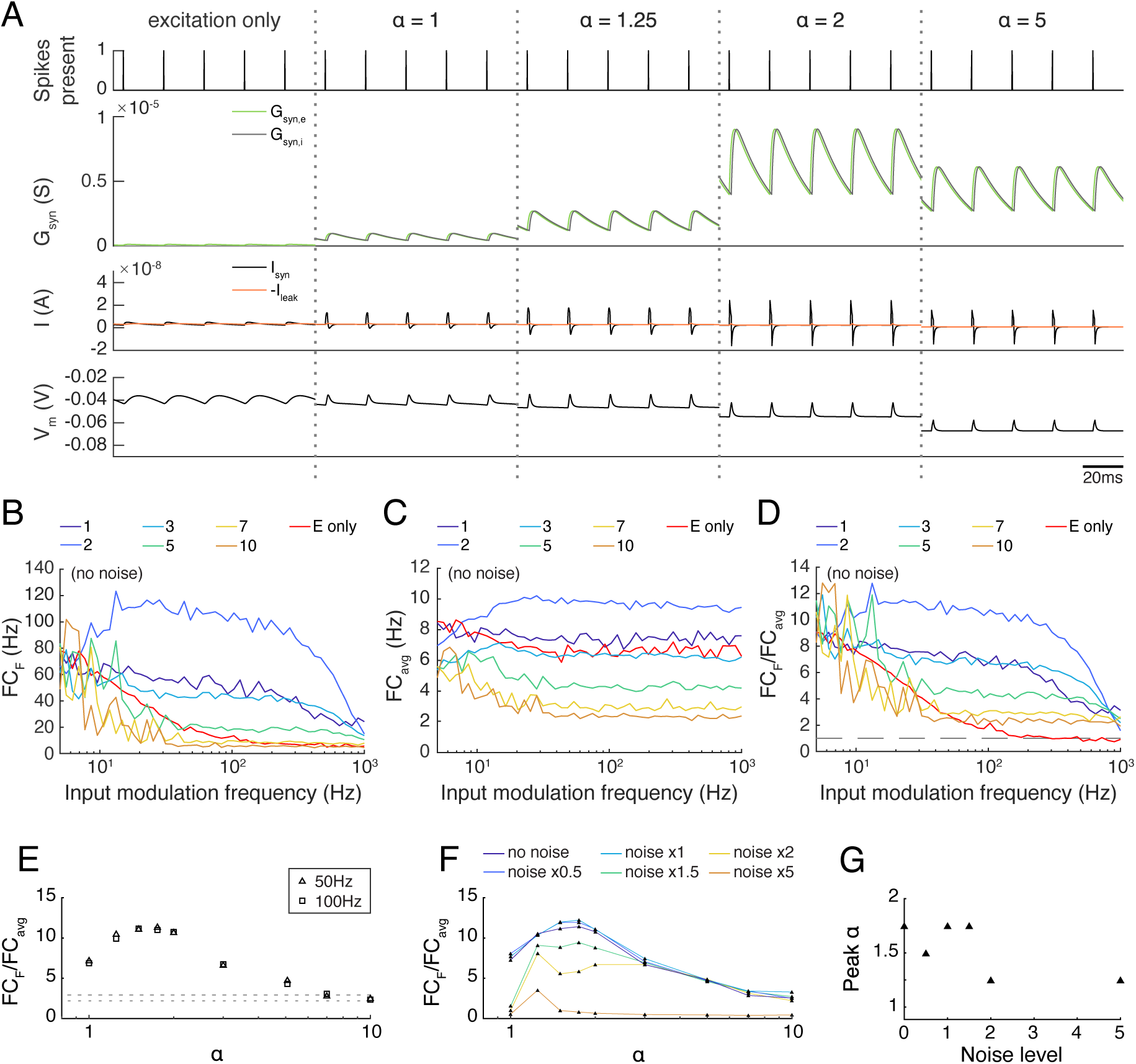
The transmission of paired feedforward E/I inputs is dependent on the strength of inhibition. **(A)** 0.1 s simulations of single-input model circuits in response to a 100Hz spike train (topmost), in which the model neuron is prevented from spiking. From left to right: feedforward E, paired feedforward E/I with inhibitory current scaled by α = 1, 1.25, 2, 5. The postsynaptic conductance, synaptic current, and membrane potential of the LGN neuron are simulated in response to the spike train input. Each simulation is shown 0.1s after the onset of spiking inputs. As the scaling on the inhibitory input increases, the inward current produced by each spike shortens in duration. **(B)** Fourier coefficient of spiking output for a range of input modulation frequencies F. Legend entries indicate the inhibitory current scaling coefficient α for each paired feedforward E/I model; “E only” indicates the feedforward E model. **(C)** Mean Fourier coefficient over all frequencies (FC_avg_). **(D)** Normalized output response over a range of frequencies F. Dashed line indicates FC_F_/FC_avg_ = 1. **(E)** The influence of scaling the inhibitory current by α on transmission FC_F_/FC_avg_, measured at F = 50Hz (triangles) and 100Hz (squares). Dashed lines indicate feedforward E model at 50Hz (top) and 100Hz (bottom). The paired feedforward E/I model exhibits greater transmission with stronger inhibition, plateauing at α = 1.5. **(F)** The influence of excitatory background noise on transmission FC_F_/FC_avg_ with different levels of inhibitory scaling α, measured at F = 50Hz. Legend entries indicate relative background current strength. **(G)** Inhibitory scaling coefficients α that produced the strongest transmission for each background noise level shown in (F). For these models, moderate inhibition (1.5 ≤ α ≤ 2) produces greater transmission when excitatory background noise is weak. As background noise becomes stronger, the model performs best with slightly scaled inhibition (α = 1.25).

The synaptic currents and postsynaptic potentials generated by excitatory-only projections were highly blurred in time and exhibited substantial temporal summation compared to those delivered by FFEI projections. In order for output spike timing to be precise, the input must be very brief in time, and synapses with NMDA cannot support this tight precision. FFEI projections delivered pulse-like current injections that were concentrated in time in a manner that varied with the total amount of inhibition α. However, sharp, high frequency inputs are highly filtered by the neuronal membrane, so the brief input to the neuron must have a very high amplitude. This can be produced by strong E and I synaptic weights, and the delay between the two serves to allow enough rapid current to produce spikes without causing a long-lasting trend in the membrane voltage. Therefore, while inputs at high frequencies are highly filtered, the large, sharp current pulses produced by the FFEI input configuration allows spiking transmission at those high frequencies despite the high filtering.

To examine the optimal amount of feed-forward inhibition, we plotted the transmission of feed-forward signals as a function of input modulation frequency (Fig. 3BCD). As before, synaptic weights were set so that transmission at an input modulation of 5 Hz was equal to about 75 Hz. Transmission at high frequencies was most efficacious when α was between 1.5 and 3. Larger values of α suppressed transmission because the level of inhibition was too high.

In many neural circuits, ongoing activity provides a background input that might influence the feed-forward transmission of information. To investigate how these background signals might influence the level of inhibition needed for optimal feed-forward transmission, we modeled this background input as 50 noisy independent Poisson inputs, with varying excitatory synaptic input strengths (Fig. 3FG). As the level of ongoing background input increased, relatively smaller weights were needed on the feed-forward inhibitory synapse for optimal transmission.

Presumably, the higher background depolarization when background noise is introduced causes a larger inhibitory current for the same conductance, and the optimal inhibitory conductance is lower; higher values would suppress transmission as they do in the no-noise case.

### Influence of inhibitory delay on feed-forward transmission

Another factor that should have a large influence on the temporal capabilities of feed-forward synapses is the delay between the feed-forward excitatory and inhibitory inputs. To examine this dependency, we varied the delay between excitation and inhibition systematically for a wide range of input modulation frequencies (Fig. 4). Shorter delays allowed transmission at much higher temporal frequencies, and a delay of 20 ms produced a transmission profile that was almost as slow as excitatory inputs alone. Therefore, a short delay between feed-forward excitation and feed-forward inhibition is necessary to achieve the high temporal frequency transmission characteristics reported here.

**Figure 4.**
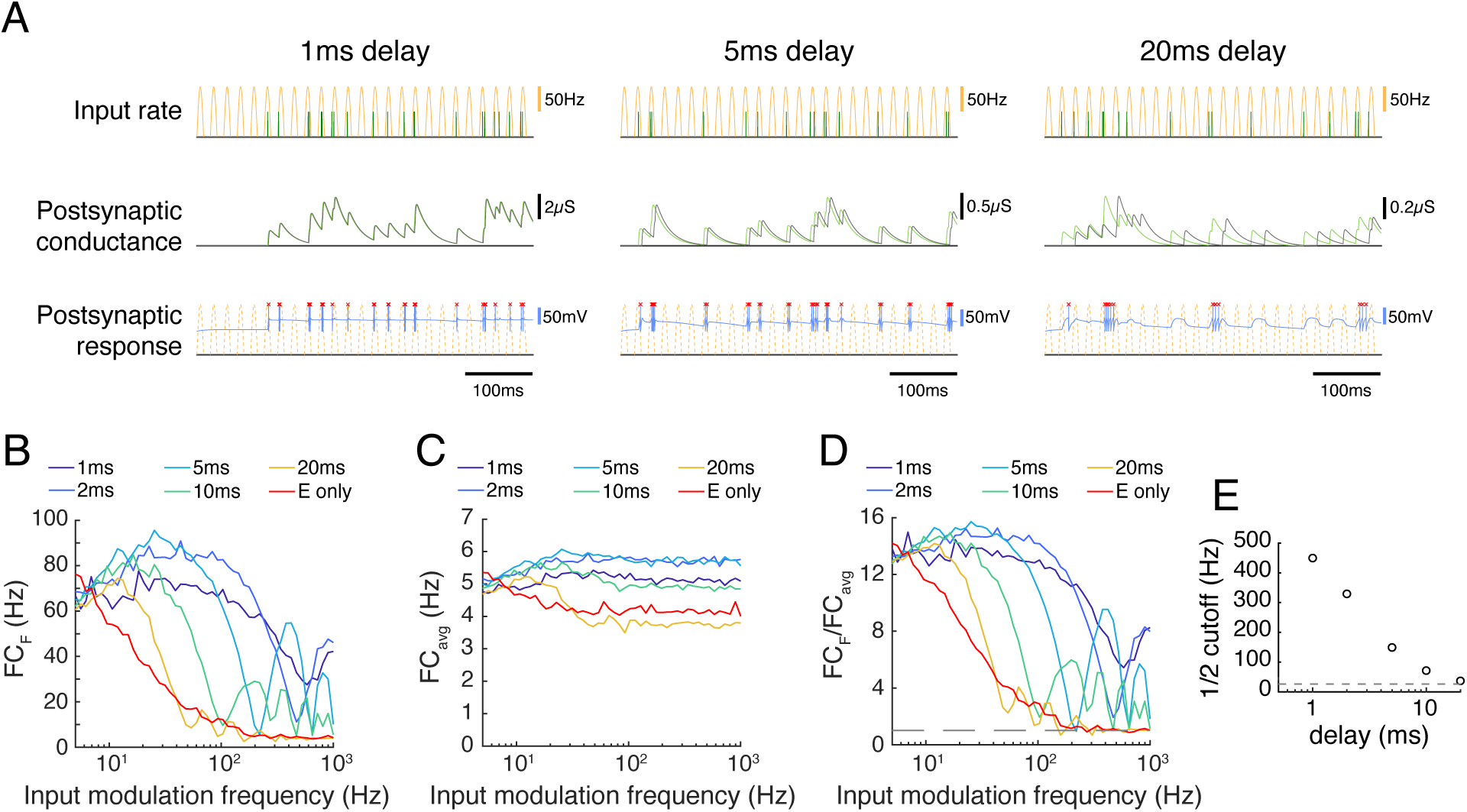
The transmission of paired feedforward inputs is dependent on the timing of inhibition. **(A)** 3 single-input model circuits, in which the LGN neuron receives feedforward E/I inputs with different inhibitory delays Δ. Each model is simulated for 0.5s with input parameters F = 50Hz, PR = 100Hz. Postsynaptic conductance (excitatory: light green, inhibitory: gray) and the membrane potential of the LGN cell (blue) are simulated in response to RG and interneuron input. LGN firing times (red x) and postsynaptic response is compared to the underlying input characteristics (dashed yellow). From left to right: Δ = 1ms, 5ms, 20ms. **(B)** Fourier coefficient of spiking output for a range of input modulation frequencies F. Legend entries indicate the inhibitory delay for each paired feedforward E/I model; “E only” indicates the feedforward E model. **(C)** Mean responses over all Fourier frequencies (FC_avg_). **(D)** The normalized output response (FC_F_/FC_avg_). Dashed line indicates FC_F_/FC_avg_ = 1. **(E)** the input modulation frequency at which FC_F_/FC_avg_ is ½(FC_F_/FC_avg_ at F = 5Hz). Dashed line indicates the ½ cutoff for the feedforward E model. With a shorter inhibitory delay, the paired feedforward E/I model reliably transmits information at higher frequencies.

### Paired feedforward excitatory and inhibitory inputs act as a clock or trigger that can organize feedforward computation across layers or networks

Neural circuits differ from conventional digital electronic circuits in that there is no external clock signal that tells neurons when they should begin evaluating their inputs (von Neumann, 1963). In digital electronics, a gate only updates its output when the clock signal indicates that it should re-scan its inputs. One classic idea suggests that neural circuits operate by attractor dynamics and therefore do not have a need for clocks (Hopfield and Tank, 1986; Miller, 2016). However, attractor dynamics take time to converge, but organisms are able to make extremely fast decisions in a few hundred milliseconds post-stimulus (Thorpe et al., 1996; Sherwin et al., 2012). A feedforward trigger would be useful for performing such fast computations. Here we examined whether paired feedforward excitation and inhibition could provide temporal organization of their own input, in essence providing the feedforward input values (“what to evaluate”) and clock timing information (“when to compute”) in a single mechanism.

We studied transmission across 4 cells in successive network levels in 3 different models. One might imagine that these cells represent the same receptive field location across these 4 network levels, so they have direct connections. However, each cell was also assumed to have local, ongoing excitatory input from its local network that was modeled as 50 noisy Poisson inputs that were not related to the feedforward input. In one model, these 4 neurons were not connected at all, so that baseline activity of these neurons could be studied as a control. In another model, these neurons were connected using FFE projections. These connections represent the simplest feedforward cross-network computation that we could imagine, which is y = x. This computation was chosen because of its ease of evaluation, but these results should extend for any convergent feedforward computation where the output is a function of the inputs at a particular time. Finally, we examined a model where cross-network transmission was provided by FFEI synapses across the 4 model network levels.

We evaluated how well the cells could transmit information forward to their subsequent levels, despite the ongoing local input that was unrelated to the feedforward inputs. We assessed the temporal fidelity of the computation using the same frequency analysis that we performed in Fig. 1. We provided an input firing rate that was modulated at a particular frequency that was varied, and examined the Fourier coefficient (Fig. 5B), overall activity (Fig. 5C), and the normalized Fourier coefficients that indicated how specifically the output response followed the input frequency (Fig. 5D). When FFE projections were used, the network level 4 output neuron did not efficaciously follow the input modulation frequency for frequencies greater than about 5 Hz. However, when FFEI projections were used, the network level 4 output neuron followed the input very well at input modulation frequencies as high as 100 Hz. This difference between FFE and FFEI was robust over a variety of feedforward synaptic strengths (Fig. 5E). Even when feedforward firing rates were set so that the output neuron was driven at 20 Hz for 5 Hz input modulation, FFEI projections provided more than an 8-fold increase in transmission compared to FFE inputs modulated at 50 Hz, and differences were more pronounced for higher synaptic strengths (Fig. 5F). The ½ cutoff frequency for the same network was more than 5-fold greater (Fig. 5G).

**Figure 5.**
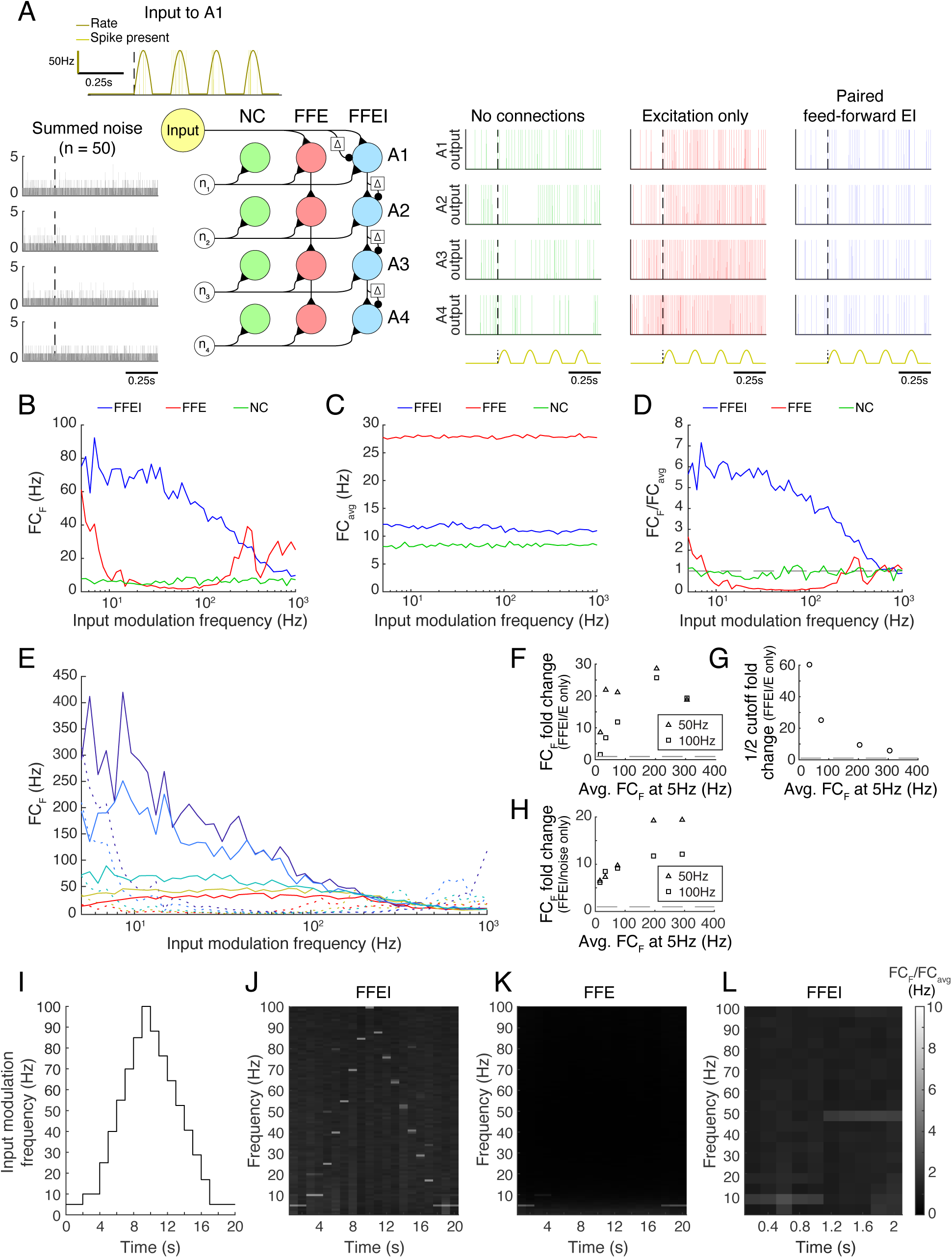
Paired feedforward E/I input allows time-locked high frequency computation across multiple brain layers or networks in the absence of an explicit external clock. **(A)** 3 model circuits exhibiting different forms of y=x transmission over 4 network levels (A1-A4): no connections between networks (NC), feedforward E-only (FFE) connections, and paired feed-forward E/I (FFEI) connections. Each circuit receives the same Poisson input (yellow) at A1, with input firing rate a rectified sine wave of frequency F and amplitude PR (dark yellow). The circuits also receive 50 “internal” noisy Poisson excitatory inputs at A1-A4 (gray, left). Simulations (NC: green, FFE: red, FFEI: blue) with input parameters F = 5Hz, PR = 100Hz shown (repeated at bottom). **(B)** For a range of input modulation frequencies F, the Fourier coefficient of the A4 output at F (FC_F_) was substantially higher for FFEI connections than for FFE connections. **(C)** Mean Fourier coefficient of the A4 output over all frequencies (FC_avg_). **(D)** Normalized Fourier output for a range of input modulation frequencies F. Dashed line indicates FC_F_/FC_avg_ = 1. FFEI allows the network levels to follow computations at higher frequencies than FFE. **(E)** Same as C, except that synaptic strength P_max_ was adjusted to produce a range of Fourier coefficient values at an input modulation frequency at 5 Hz (weak to strong: red to dark blue). Solid lines indicate the feedforward E/I model, and dashed lines indicate the feedforward E model. **(F)** Summary data from E, showing fold change advantage of FFEI over FFE at F = 50 Hz (triangles) and F = 100 Hz (squares) for various synaptic strengths (FC_F_ at 5 Hz shown on x axis). Dashed line indicates 1. **(G)** Same, but fold change in the ½ cutoff shown. The ½ cutoff fold change could not be measured for the weakest FFEI/FFE pair in **(E)**, as the paired feedforward E/I model does not have a ½ cutoff within the range of considered F values. **(H)** For each paired feedforward E/I model shown in **(E)**, the FC_F_ fold change with respect to the model with no connections (noise only) is shown. Dashed line indicates a fold change of 1. **(I)** Over a 20 second trial, a rectified sinusoidal Poisson input with a changing input modulation frequency was provided to the circuits exhibiting FFE and FFEI connections. **(J)** The normalized Fourier spectrum of the A4 output with FFEI connections, averaged over 1 second bins. Spectrum is averaged over 50 trials. **(K)** Same as **(J)**, but with FFE connections, which cannot follow high frequencies. **(L)** The FFEI circuit was provided with a rectified sinusoidal Poisson input with an input modulation frequency that was stepped from 10 Hz to 50 Hz at t = 1 s. The Fourier spectrum of the A4 output is averaged over 0.2 second bins, showing a rapid transition of the output firing.

Interestingly, when FFE projections were used, the Fourier coefficient of the network level 4 output peaked several times at higher input frequencies (300-1000 Hz). This effect can likely be credited to over-excitation of the network level 4 neuron, as seen in Fig. 5A. Indeed, at most frequencies, the network level 4 neuron with FFE is likely firing at its maximum rate, due to the amplification of excitatory input over the circuit. As a result, high FC measurements at high input frequencies cannot necessarily be attributed to the circuit following the inputs, but may instead result from the physical features of the neuron allowing for maximal firing at these rates. In either case, the Fourier coefficient peaks at these high input frequencies do not stand out from the overall power of the output response, as shown in Fig. 5D.

One may ask how much of these frequency responses could have arisen by chance due to the random activity that is provided to each network level. The fold increase in Fourier coefficient responses at the input modulation frequency is shown in Fig 5H. Even at the lowest synaptic strength tested, FFEI projections provided responses that were at least 5 times those of the noise-only network, and this difference greatly increased when input strength was increased.

So far, we have only investigated how inputs of fixed modulation frequency are transmitted over multiple network levels. While this addresses how different network levels can exhibit synchronized activity, it does not directly confront how these networks can adapt to changing inputs. We therefore constructed a rectified sinusoidal Poisson input with a changing modulation frequency over time (Fig. 5I). After changing the input modulation frequency, we held the input modulation frequency constant for 2 seconds in order to produce a reliable estimate of the Fourier coefficients. We then provided this input to the FFE and FFEI models and examined the normalized FC spectrum over time (Fig. 5JK), which measured how closely the network level 4 output followed a range of frequencies over the course of the input. With FFE projections, the network level 4 output only noticeably followed the stepping input at low frequencies (5-10Hz). Meanwhile, with FFEI projections, the network level 4 output followed the entire stepping input fairly well. Furthermore, the network level 4 output did not exhibit significant residual power at frequencies represented earlier in the input, suggesting that the FFEI model was capable of quickly adapting to the changing input frequency. To more closely examine the speed at which the FFEI model transitions between inputs of different modulation frequencies, we provided the model with a Poisson input that increased from F = 10 Hz to F = 50 Hz at time = 1 second, and computed the FC spectrum of the network level 4 output over smaller time intervals (0.2 seconds) (Fig. 5L). With this increased temporal resolution, the FFEI model rapidly adapts to an increase in the input frequency.

Taken together, these results indicate that FFEI projections organize multi-level, hierarchical computations that are also being driven by ongoing incidental activity. Further, FFEI projections allow these computations to occur with high fidelity even at high temporal frequencies. In this way, FFEI projections serve to deliver both a signal as well as the timing signal necessary to organize feedforward computation across layers of processing.

### Paired inputs of opposite sign can organize computation in neuromorphic circuits

We have already noted that paired feedforward excitation with slightly delayed inhibition is a common circuit motif in the brain, but the concept can be used more generally for computations with either sign. Although we are not presently aware of feedforward inhibition that is followed by delayed excitation in a neural circuit, this motif can be used in artificial circuits to organize feedforward computations at high speeds that exceed the time constant of integration of the individual elements.

We demonstrate the use of mixed feedforward inputs in a model LIF circuit that performs the exclusive-or (XOR) computation of 2 inputs (Fig. 6). In this computation, the output neuron should exhibit a response if either of the 2 inputs is positive, but it should be silent if there is no input or if both inputs are active. To transmit a positive postsynaptic signal, we used FFEI synapses employing excitation that was followed by identical but delayed inhibition (Fig. 6A). To transmit a negative signal, we employed inhibition that was followed by identical but delayed excitation. Input weights for cells in the first layer were set so that one input was positive and the other input was negative. Further, the input with a weight that was positive in the first cell had a negative weight in the second cell. The neuron in the second layer responded if either of the cells in the first layer responded.

**Figure 6.**
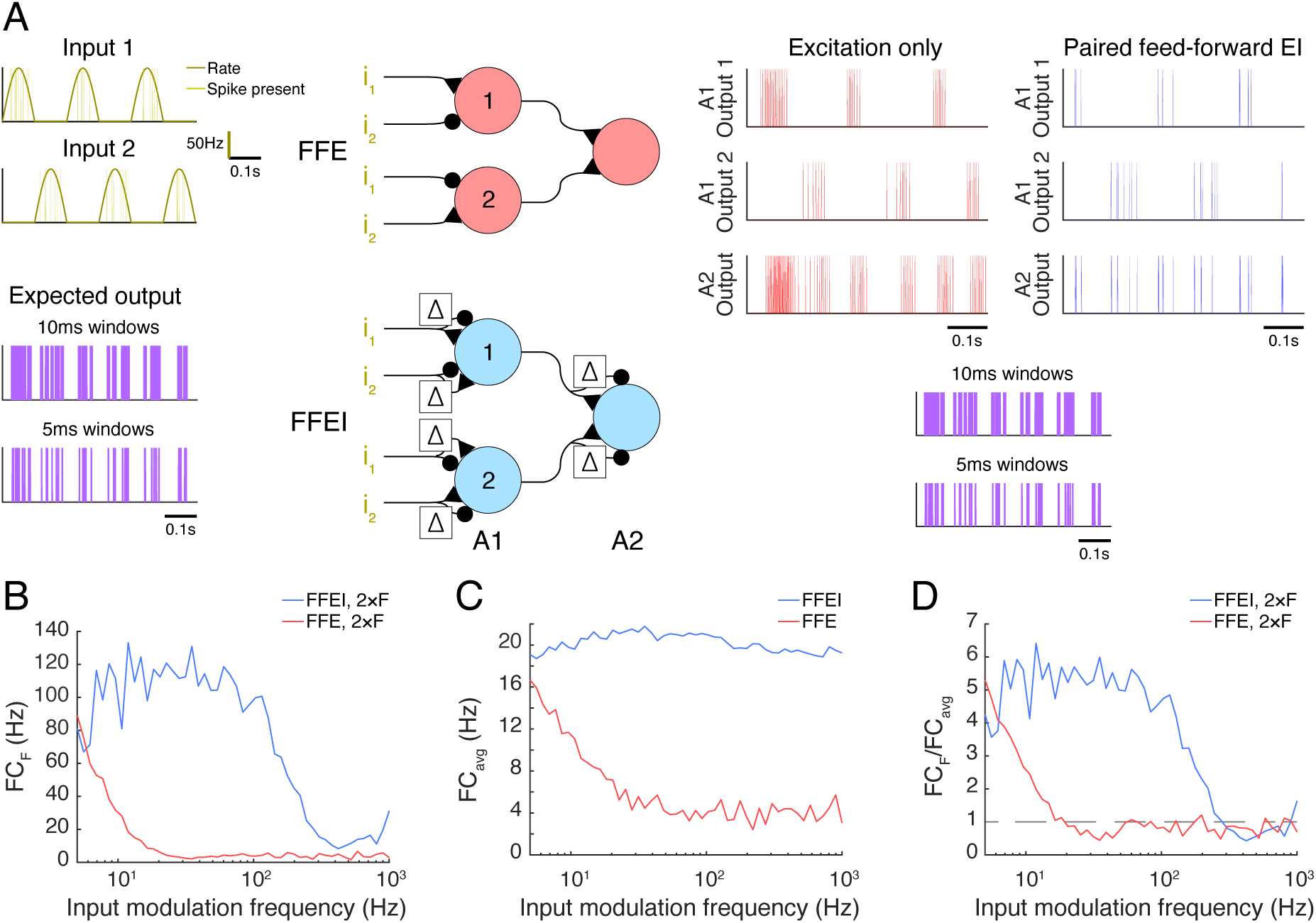
Paired feedforward E/I allows high frequency computation in artificial neuromorphic circuits. **(A)** Model circuits utilizing either feedforward single-sign inputs only (center, top) or paired mixed-sign inputs (center, bottom) to perform an exclusive-or (XOR) computation. The cell in layer A2 fires only when either Poisson input i_1_ or i_2_ (dark yellow) is active, but not both. A 0.6s simulation of each model circuit with inputs driven out-of-phase (π) is shown with input parameters F = 5Hz, PR = 100Hz. The out-of-phase input should drive the XOR circuit strongly at twice the input modulation frequency, because one input is firing when the other is not at two phases in the input wave.The simulated output spikes of cells 1 and 2 in A1 and the cell in A2 are shown to the right for each model circuit (single-sign: red, mixed-sign: blue). The expected output of the XOR computation between i_1_ and i_2_ (purple), calculated in 10ms or 5ms windows, is shown below for comparison with A2 outputs. **(B)** A2 output Fourier coefficient at 2xF (FC_F_) for a range of input modulation frequencies F. **(C)** A2 output mean Fourier coefficient. **(D)** A2 output normalized Fourier coefficient. Dashed line indicates FC_F_/FC_avg_ = 1.

Once again, we provided input at different modulation frequencies, although we shifted the two inputs so that they were 180° out of phase (Fig. 6A). Therefore, the XOR output should be produced at twice the input frequency (the cell should respond strongly when either input is at its positive phase, and weakly or not at all when the inputs are turned off). Once again, we found that the circuit with mixed FFEI projections could follow the XOR computation over high frequencies, while a comparison FFE circuit fell off rapidly with frequency, for both the raw response (Fig. 6B) and for the fraction of the total response (Fig. 6C) that was at the stimulus frequency Fig. 6D). We compared another input case (with the 2 inputs shifted by 90°) in Supplementary Figure 6-1.

We conclude that the concept of mirroring a feedforward input with a delayed negative copy can organize high frequency computations across multi-layer neuromorphic circuits in much the same way that a clock organizes computations across digital electronic circuits.

### In principle, AMPA-only currents allow high frequency computations

Most excitatory synapses in the mammalian brain employ a combination of the AMPA and NMDA receptors. AMPA receptors respond rapidly and briefly to neurotransmitter, while NMDA receptors remain open for longer periods of time (50-150 ms). However, there are a few instances of excitatory synapses that primarily involve AMPA receptors. One example is the calyx of Held synapse in the mature mammalian auditory brain stem; notably, this region often processes high frequency temporal information in the form of auditory input (Nakamura and Cramer, 2011). This raises the question of whether neural circuits could, in principle, achieve the same fast computation as FFEI by using feedforward inputs that were comprised entirely of fast AMPA channels.

To address this question, we compared feedforward transmission in synapses that were comprised of AMPA receptors only, AMPA and NMDA receptors only (like FFE), and AMPA, NMDA, and GABA receptors (like FFEI) in Figure 7. We again provided rectified sinusoidal input at an input modulation frequency, and examined the Fourier coefficient of the response at the input frequency (Fig. 7B), the average response across all frequencies (Fig. 7C), and the normalized output frequency (Fig. 7D). Unlike in previous simulations with feedforward excitatory transmission alone, accurate high-frequency transmission was possible when only fast-closing AMPA channels were present. However, when slow-closing NMDA channels were added, the excitatory conductance and the spiking response became blurred, leading to poor transmission of high-frequency information. High-frequency transmission was restored by inhibition via GABA channels.

**Figure 7.**
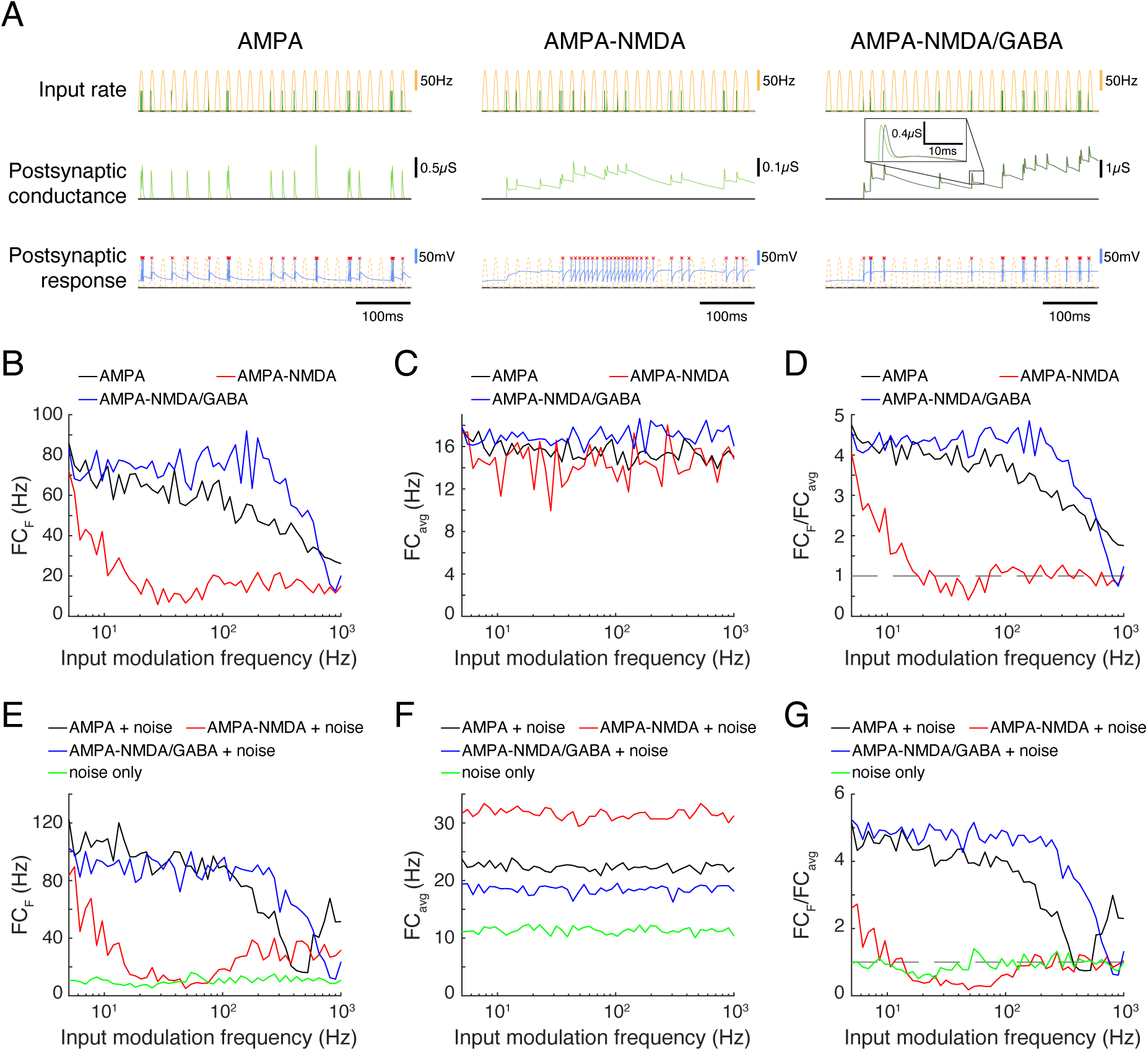
Feedforward excitatory input that is restricted to AMPA channels allows high frequency computation but is less robust to noise compared to FFEI inputs. **(A)** 3 triad synapse-like models, in which the LGN neuron receives inputs via different sets of synaptic channels. Each model is simulated for 0.5s with input parameters F = 50Hz, PR = 100Hz. Postsynaptic conductance (excitatory: light green, inhibitory: gray) and the membrane potential of the LGN cell (blue) are simulated in response to RG and interneuron input. LGN firing times (red x) and postsynaptic response is compared to the underlying input characteristics (dashed yellow). Left: inputs are transmitted via AMPA channels only. Center: inputs are transmitted via AMPA and NMDA channels. Right: inputs are transmitted via AMPA, NMDA, and GABA-like channels. **(B)** Fourier coefficients (FC_F_) for a range of input modulation frequencies F. **(C)** Fourier coefficients averaged over all response frequencies (FC_avg_) for a range of input modulation frequencies F. **(D)** Normalized Fourier coefficients (FC_F_/FC_avg_). Dashed line indicates FC_F_/FC_avg_ = 1. AMPA-only synapses, with their shorter time constants, allow transmission at higher frequencies than synapses with both AMPA and NMDA, and are comparable to AMPA-NMDA-GABA-like synapses for frequencies less than about 20 Hz. **(E)** In addition to the rectified sinusoidal Poisson input, the LGN cell receives 50 noisy Poisson excitatory inputs. Fourier coefficients (FC_F_) are shown for a range of input modulation frequencies F; “noise only” denotes the absence of rectified sinusoidal input. **(F)** Same as **(C)**, but in the presence of LGN noise. **(G)** Same as **(D)**, but in the presence of LGN noise. FFEI slightly enhances transmission in the presence of noise compared to FFE with AMPA synapses alone.

To investigate the effects of local background activity on the ability of the synapses to follow high-frequency inputs, we provided 50 noisy Poisson inputs, modeled as in Fig. 5, and examined the output response (Fig. 7EFG). In the presence of excitatory noise, the FFEI synapse overall followed rectified sinusoidal inputs more effectively than the AMPA-only feedforward synapse. These observations can be attributed to the overall output power, which was somewhat higher for the AMPA-only feedforward synapse than for the FFEI synapse (Fig. 7F).

These results suggest that AMPA-only feedforward synapses provide some of the benefits of FFEI with respect to high frequency computation. However, FFEI synapses are more effective than AMPA-only feedforward synapses when noisy inputs are also provided, suggesting that inhibitory GABA channels may play a role in filtering out excitatory noise and thus overriding ongoing computations.

### High frequency transmission in a triad synapse model with synaptic depression

Up to now, we have not considered an important feature of synaptic transmission in real triad synapses in the LGN: short-term dynamics (Abbott et al., 1997; Tsodyks and Markram, 1997; Varela et al., 1997; Carandini et al., 2002; Swadlow et al., 2002; Boudreau and Ferster, 2005; Higley and Contreras, 2006). The retino-geniculate excitatory and inhibitory synapses, as measured *ex vivo* in mouse, both exhibit substantial short-term depression (Eysel, 1976; Koch, 1985; Chen and Regehr, 1999; Blitz and Regehr, 2003, 2005). We developed a depression model for excitatory and inhibitory synapses based on experimental data from mouse (Chen and Regehr, 1999; Blitz and Regehr, 2003, 2005) using depression equations from Varela et al. (1997), and added AMPA, NMDA, GABA_A_ and GABA_B_ currents (Supplementary Figure 8-1). NMDA and GABA_B_ currents have long time courses, and it is necessary to have compatible time courses for the paired feed-forward input to allow high frequency transmission.

Adding depression dynamics also meant that we needed to consider the state of the synapses at the beginning of the simulation. *In vivo* in the cat, retinal ganglion cells exhibit relatively high spontaneous firing rates of tens of spikes/sec (Kuffler, 1953), so it is unlikely that retinal ganglion cell synapses would commonly be in a completely undepressed state; the neuron would need to be shown a stimulus that suppresses its firing, such as showing an ON spot to an OFF-center neuron, to achieve such a state. Therefore, although we began our simulations with the synapses in a completely undepressed state, we did not analyze the output until after 1 second had elapsed.

Representative simulations for AMPA + NMDA (FEI) and AMPA, NMDA, GABA_A_, and GABA_B_ (FFEI) synapses are shown in Figure 8AB. When the simulations begin, there is strong excitatory transmission, but excitation and inhibition become more balanced as the simulation runs. Synaptic weights were chosen for excitatory and inhibitory synapses so as to allow moderate firing in the FFEI case after 1 second of simulation; inhibitory inputs were removed for the FFE case, and the excitatory weights were adjusted so that the firing rate for inputs at 5 Hz modulation exhibited 70 Hz after 1 second of simulation, matching the FFEI simulations. With synaptic depression, transmission (Figure 8CDE) is improved for FEI synapses compared to the simulations without synaptic depression, consistent with previous theoretical analyses (Abbott et al., 1997). However, transmission at higher temporal frequencies is better with the FFEI synaptic arrangement as compared to FEI.

**Figure 8.**
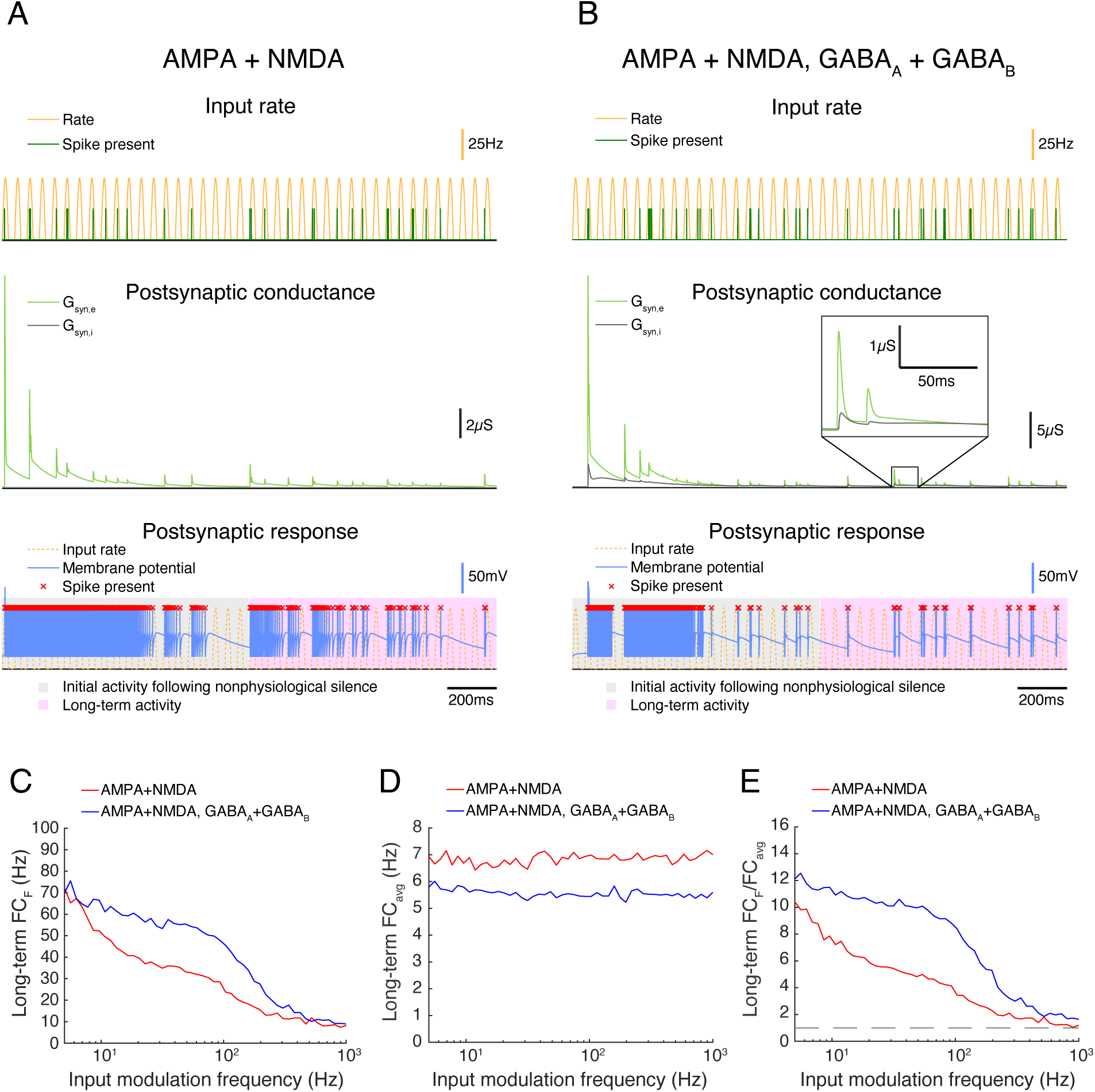
In triad synapse models exhibiting synaptic depression, paired feedforward E/I enhances high-frequency transmission. **(A-B)** 2 triad synapse models, in which the LGN neuron receives inputs via different sets of channels exhibiting synaptic depression. Each model is simulated for 2s with input parameters F = 20Hz, PR = 50Hz. Postsynaptic conductance (excitatory: light green, inhibitory: gray) and the membrane potential of the LGN cell (blue) are simulated in response to RG and interneuron input. LGN firing times (red x) and postsynaptic response is compared to the underlying input characteristics (dashed yellow). The output response is divided into 2 components over time: a 1s period of initial activity following nonphysiological silence (gray highlight) and the subsequent long-term activity (pink highlight). **(A)** Inputs are transmitted via AMPA and NMDA channels only. **(B)** Inputs are transmitted via AMPA, NMDA, GABA_A_, and GABA_B_ channels. **(C)** Fourier coefficients (FC_F_) for a range of input modulation frequencies F. Models are simulated for 5s, and Fourier coefficients are computed 1s after the start of each simulation to examine the long-term activity. **(D)** Fourier coefficients averaged over all response frequencies (FC_avg_) for a range of input modulation frequencies F. **(E)** Normalized Fourier coefficients (FC_F_/FC_avg_). Dashed line indicates FC_F_/FC_avg_ = 1. With synaptic depression, AMPA-NMDA-GABA synapses allow for transmission at higher frequencies than synapses exhibiting excitation alone.

We modulated the input rate sinusoidally from 0 to 50 spikes/sec, a reduction from the 0 to 100 spikes/sec we used in the prior examples. Blitz and Regehr (2005) noted that high frequency spike trains (50-100 Hz) caused such substantial depression that the inhibitory synapse effectively stopped transmitting. Here, it is important to note that we are modulating the firing rate probability of the input sinusoidally, and a 50 Hz peak firing rate probability modulated sinusoidally will rarely result in spikes that have an inter-spike-interval of 20 ms (1/50 Hz). Visual firing rates are typically quantified in a trial-averaged fashion and expressed as the probability of firing in a small time bin, and we are generating spikes according to such a process. At sustained high frequency input rates >50Hz, it is likely that transmission of both excitation and especially inhibition would be very reduced, although it is unclear how common this situation of sustained high input rates would be *in vivo*, or if synaptic depression is altered *in vivo* or across species.

### Transmission is limited by inhibitory neuron time constants when FFEI is provided at the circuit level

While the FFEI motif is sometimes found in inputs onto individual neurons, such as in the retinogeniculate triad synapse, the FFEI motif is also commonly found in inputs to circuits such as in the hippocampus (Buzsaki, 1984; Pouille and Scanziani, 2001; Bhatia et al., 2019) or the cerebral cortex, both in inputs from thalamus (Agmon and Connors, 1991; Swadlow, 2003) and interareal projections (Yang et al., 2013). In these networks, excitatory projections from other areas make connections with separate populations of excitatory and inhibitory neurons, and the targeted inhibitory neurons provide rapid but slightly delayed inhibition onto the recipient excitatory neurons. Further, the excitatory and inhibitory neurons typically receive converging inputs from many cells, so that in principle the excitatory and inhibitory inputs to the local circuit may not be entirely matched. That is, the strong 1-to-1 relationship between each feedforward excitatory postsynaptic potential and each feedforward inhibitory postsynaptic potential that is found at the triad synapse is not present in circuit-level projections; instead, feedforward excitation and inhibition are more loosely related.

Before we begin studying how FFEI can impact temporal computations at the circuit level, it is worth pausing to consider whether all cortical circuits might take advantage of any properties that we uncover. In the thalamus, it is clear that visual, somatosensory, and auditory neurons can follow sensory inputs at high frequencies. However, in visual (Hawken et al., 1996), somatosensory (Chung et al., 2002), and auditory cortex (Creutzfeldt et al., 1980; Wehr and Zador, 2005), cortical neurons primarily respond to stimulation at much lower stimulus repetition frequencies (<20 Hz) than do thalamic neurons. That is, primary sensory cortex does not seem to segment responses to sensory stimulation at frequencies much higher than 20 Hz. However, in both sensory and higher cortical areas and in the hippocampal formation, 40 Hz activity (gamma activity) is prominent (Fries et al., 2007) and can be induced by driving feedforward interneurons (Cardin et al., 2009). It has been suggested that in hippocampal circuits, each gamma cycle might encode representations of individual items (Lisman and Idiart, 1995; Fries et al., 2007; Lisman and Jensen, 2013), and 40 Hz sensory stimulation has been reported to be protective against neurodegeneration (Iaccarino et al., 2016). So we feel it is worth considering whether circuit-level FFEI input can allow circuits to follow inputs at such high frequencies.

Here, we examined a model network with 100 feedforward excitatory inputs that arrived at an LIF excitatory cell and an LIF inhibitory cell (Fig. 9A). Following our previous methods, the firing rate of each input was modulated sinusoidally at an input modulation frequency. In this model, all inputs were driven at the same phase (Fig. 9B). We examined the Fourier coefficient of the output responses at the input modulation frequency (Fig. 9C), the average response across all frequencies (Fig. 9D), and the ratio of these quantities (Fig. 9E).

**Figure 9.**
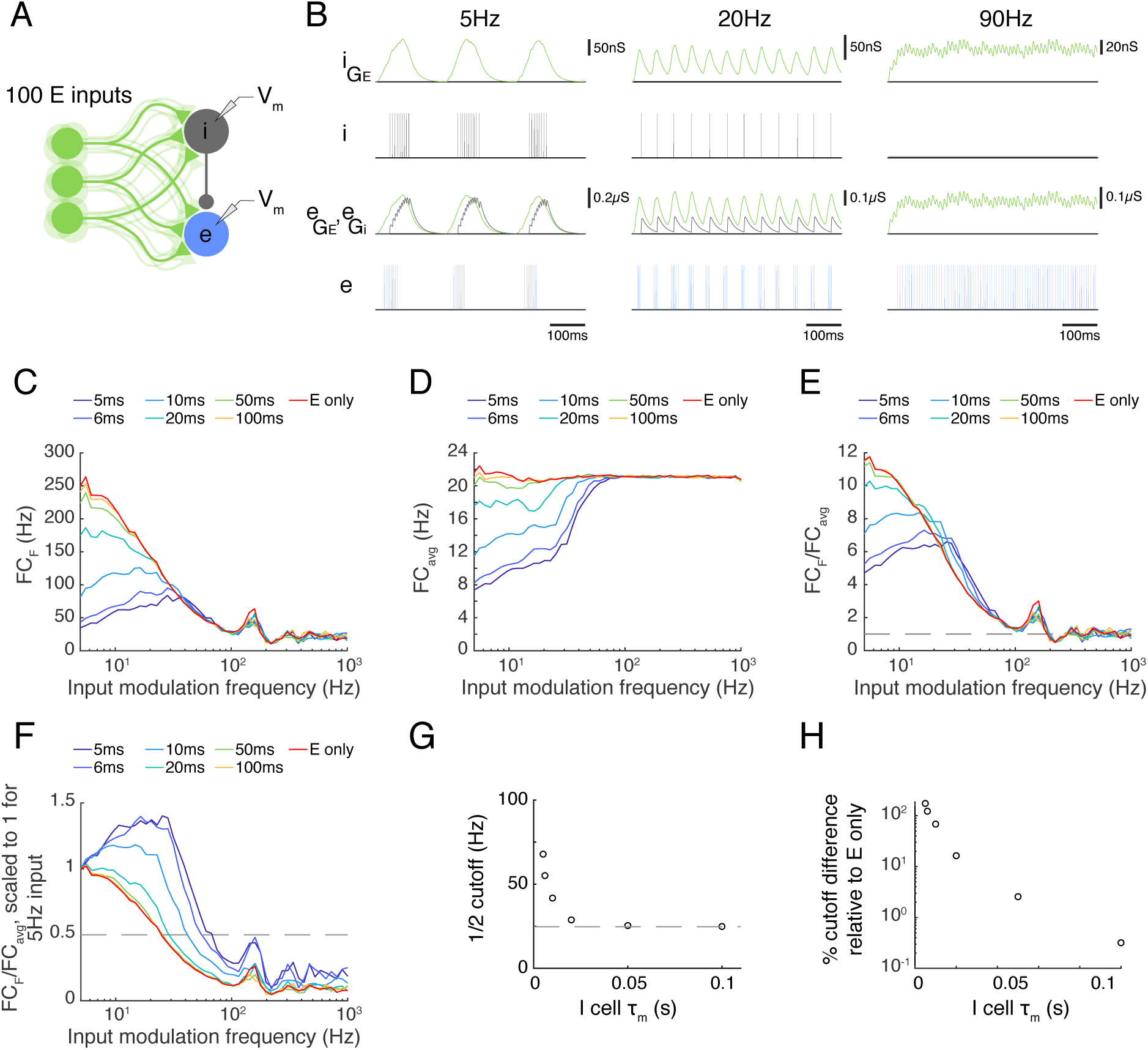
Paired feedforward E/I in a multi-input model allows computations that are frequency-limited by the membrane time constant of the feedforward inhibitory neuron. **(A)** A paired feedforward E/I model circuit with 100 input cells (light green) providing excitatory (E) input to a cortical neuron e (blue) and to an interneuron i (gray), which in turn provides inhibitory input to the cortical neuron. **(B)** 0.6s simulations of the model circuit with Poisson input parameters F = 5Hz (left), 20Hz (center), and 90Hz (right), and PR = 100Hz. The excitatory postsynaptic conductance (light green) in the i (i_GE_) and e cells (e_GE_) are simulated in response to the summed input. This excitatory input produces spiking of the i cell (dark gray), resulting in an inhibitory postsynaptic conductance in the e cell (light gray, e_GI_). The combination of excitatory and inhibitory inputs produces the observed spiking in the e cell (blue). Notably, there is an absence of inhibitory input to the e cell at 90Hz. **(C)** Fourier coefficient at input frequency F (FC_F_) for a range of input modulation frequencies. Legend entries indicate the value of the interneuron membrane time constant τ_m,I_ for each paired feedforward E/I model; “E-only” indicates a feedforward E model, in which the interneuron is not present. **(D)** Fourier coefficients averaged over all response frequencies (FC_avg_) for a range of input modulation frequencies F. **(E)** Normalized Fourier coefficients (FC_F_/FC_avg_). Dashed line indicates FC_F_/FC_avg_ = 1. **(F)** Scaled FC_F_/FC_avg_ values from (E) such that FC_F_/FC_avg_ = 1 for F = 5Hz. The dashed line indicates FC_F_/FC_avg_ = 0.5, and intersects with each FC_F_/FC_avg_ curve at the ½ cutoff. **(G)** The τ_m,I_ dependence of the ½ cutoff. Dashed line indicates the ½ cutoff for the feedforward E model. **(H)** Percent difference in the ½ cutoff relative to the feedforward E model as a function of τ_m,I_. Circuit-level feedforward E/I input allows excitatory neurons to follow higher temporal frequencies than those receiving feedforward excitatory input alone, but the cut off frequency values are smaller than for the triad synapse, where the interneuron cell body integration does not intervene.

The circuit-level FFEI exhibited some similarities to and differences from the single FFEI input situation. One difference is that the circuit’s I neuron receives excitatory input without a delayed inhibitory copy, and acts just like a neuron receiving FFE input. Therefore, the frequency responses of the inhibitory neuron are the same as the E-only case in Fig. 9CDE. If we look at these responses, we see that they fall off much like FFE input to the triad synapse: the response is much attenuated at about 25 Hz, for example. At input modulation frequencies greater than this, the I neuron will not respond strongly to the excitatory input (Fig. 9B**, right**), so the I neuron’s contribution to the circuit is not present at high frequencies, and the circuit behaves like an FFE circuit. This means that the circuit-level FFEI projection has a much lower cut-off response frequency than a similarly-endowed triad-like FFEI projection.

We suspected that the membrane time constant of the inhibitory neuron would be a key parameter in determining the cut-off frequency for the circuit-level FFEI projection, so we compared responses in models where this parameter was varied. In this circuit network, the level of response output is very sensitive to the synaptic strengths of the inputs as well as the strength of the synapse from the inhibitory neuron onto the excitatory neuron. Because there were 2 non-linear ways to tune the responses to a particular input rate, we simply left the synaptic strengths unchanged while we altered the membrane time constant of the inhibitory neuron so that the models could be compared on the basis of that parameter.

In Fig. 9F, we show the normalized Fourier coefficient for these models but with the data scaled so that the output response for an input modulation of 5 Hz is equal to 1. Here it is apparent that the frequency response depends strongly on the inhibitory membrane time constant, with low values corresponding to high transmission at high input modulation frequencies. When the inhibitory time constant was 10 ms, which is consistent with experimental studies of cortical interneurons (Economo and White, 2012), the ½ frequency cut off was 40 Hz, indicating that the excitatory cell could follow input responses very effectively up to that frequency (Fig. 9G). This ½ cutoff frequency was 70% greater than the ½ cutoff frequency for a circuit with FFE projections alone (Fig. 9H).

## Discussion

We have shown that feedforward excitation and delayed inhibition can enhance the transmission of high-frequency temporal information through neural circuits, and that this enhancement occurs for a broad range of synaptic and membrane parameters. Similar to the triad synapses in retinogeniculate projections, FFEI models with low input convergence and tightly paired E/I inputs exhibit robust temporal transmission over a wide range of frequencies up to ∼100 Hz. Meanwhile, in circuits modeled after hippocampal and cortical projections, FFEI enhances high frequency transmission up to about 40 Hz, despite high input convergence (N = 100) and a looser association between excitatory and inhibitory inputs. In circuit models of hierarchical networks, we found that FFEI projections reliably transmit high frequency signals and organize activity across network levels over time, suppressing the influence of ongoing activity and essentially acting as a feedforward clock trigger. Thus, FFEI projections can act as a hybrid input, conveying both information about input value and imposing a clock-like trigger to initiate rapid computation.

### FFEI allows computation at high temporal frequencies that “break the limit” of the membrane time constant

Excitation with delayed inhibition produces a narrow time window during which spiking can occur. As a result, FFEI inputs can enhance the temporal fidelity of neural circuit outputs, as shown in experiments in hippocampal CA1 (Pouille and Scanziani, 2001; Bhatia et al., 2019) and at the retino-geniculate synapse (Blitz and Regehr, 2005). Given the high temporal precision of networks containing FFEI projections, it follows that such networks would accurately transmit high-frequency temporal information, as well. This effect can largely be attributed to the narrow spiking time window generated by paired E/I inputs, as previously described (Pouille and Scanziani, 2001; Wehr and Zador, 2003; Blitz and Regehr, 2005; Mittmann et al., 2005; Higley and Contreras, 2006; Cardin, 2018). Here we show that the residual inhibition following each input spike does not suppress the cell for long, and the cell is quickly ready to respond to subsequent FFEI inputs (Figure 3A).

Networks that propagate signals with high temporal fidelity have been called synfire chains (Abeles, 1991). We have shown here that excitatory-only feed-forward networks can produce reliable computation only at relatively low temporal frequencies. Excitatory-only feed-forward networks can reliably and quickly transmit firing rate information if each area receives ongoing balanced excitatory and inhibitory internal noisy input (Shadlen and Newsome, 1998; Chance et al., 2002; van Rossum et al., 2002; Murphy and Miller, 2009) or has detailed or loosely balanced strongly recurrent connections (Tsodyks and Sejnowski, 1995; van Vreeswijk and Sompolinsky, 1996, 1998; Hennequin et al., 2017) or balanced connectivity across cortical areas (Joglekar et al., 2018). Here we have shown that this transmission is possible with FFEI projections even if the random local input that a local area experiences or the internal connectivity of the network is not balanced: transmission at high temporal frequencies can be accomplished with balanced feedforward input alone.

It should be noted that some brain pathways use different strategies to transmit high frequency information. In the auditory brainstem, neurons respond rapidly to a transient input (Schnupp and Carr, 2009), which can be explained by the composition of the intrinsic ionic currents in the cell that allow them to act as “slope detectors” (Meng et al., 2012), although they do not respond tonically to constant current, as LGN neurons do (Suzuki and Rogawski, 1989).

The high transmission functions of FFEI could act in concert with another proposed function for the triad synapse: a state-dependent “veto” (Koch, 1985). Under this idea, the synapse would act as described here when operating in a feedforward mode, but the inhibitory neuron could be activated independently in order to suppress feedforward excitatory input to support functions such as reduced attention or internal processing. A related feedforward excitatory/inhibitory gate model was demonstrated by Kremkow et al. (2010a). In recurrent, balanced cortical networks, inhibitory synaptic strengths (Vogels and Abbott, 2009) or the correlation between excitatory and inhibitory signals can be used to gate the propagation of tonic activity through downstream networks, and this can be combined with temporal gating (Kremkow et al., 2010a). Learning rules (Vogels et al., 2011; Kleberg et al., 2014) could tune inhibition for ideal propagation at high temporal frequencies (see Figure 3).

### FFEI as a feedforward “clock” trigger mechanism

The absence of a synchronizing clock or trigger signal to organize the activity of neurons would seem to imply that neuronal circuits would be limited to performing computations that do not require such a trigger, such as attractor networks or integrators (Hopfield, 1982; Seung et al., 2000; Miller et al., 2003; Miller, 2016). FFEI acts much like a feed-forward synchronizing trigger that forces computation to occur within a short window (Buzsaki, 1984; Agmon and Connors, 1991; Swadlow and Gusev, 2000; Porter et al., 2001; Swadlow, 2003; Rock and Apicella, 2015; Bhatia et al., 2019).

Here we showed that FFEI, when used in projections across brain areas or layers, can organize computation across these regions, even when these regions are experiencing ongoing local activity (Fig. 5). Unlike a digital computer, where the clock trigger runs at a fixed rate, FFEI allows the trigger signal to be generated at any time (Fig. 5IJK) and at many rates, up to 100 Hz for triad-like synapses and up to about 40 Hz for circuit-level FFEI inputs. In low-level areas, these triggers would be generated when sensory input arrives, and computation could proceed at the speed of the sensory input. In higher-level areas, computation across two areas could proceed whenever the input area generates output, and could underlie coordination of high frequency oscillations across areas (Lisman and Idiart, 1995; Lisman and Jensen, 2013).

The degree to which high frequency information actually travels throughout the brain varies depending upon area and channel, and has been well studied in the visual system. In recordings that examined velocity tuning to single bar sweeps in both retina and LGN of X and Y cells in the cat, peak velocity tuning was slightly lower in LGN for X cells, but very comparable in retina and LGN for Y cells, with preferred velocities that ranged from about 20°/s to 100°/s (Frishman et al., 1983), indicating that spike timing information can be precisely transmitted. Consistent with this idea, retinal inputs activate LGN cells with high efficacy (Usrey et al., 1998), indicating that LGN neurons follow retinal ganglion inputs with high reliability.

Evidence from sensory cortex, however, suggests that the thalamocortical connection does not take full advantage of the potential FFEI speeds. There is a drop in peak temporal frequency tuning between LGN and cortex, with most LGN cells exhibiting strong firing for many frequencies, including those greater than 16 Hz, while few cortical neurons exhibit strong firing at 16 Hz or greater (Hawken et al., 1996; Van Hooser et al., 2003; Heimel et al., 2005; Van Hooser et al., 2013). If FFEI were the only factor, we would predict that transmission should be quite robust at 20 Hz (Fig. 9), but visual cortex attenuates these frequencies quite strongly. This drop also occurs in the somatosensory (Ahissar et al., 2001; Chung et al., 2002) and auditory systems (Creutzfeldt et al., 1980). Synaptic depression or other cortical circuit mechanisms may underlie this filtering (Chung et al., 2002; Swadlow, 2002; Boudreau and Ferster, 2005; Swadlow et al., 2005; Wehr and Zador, 2005).

Nevertheless, many cortical regions exhibit strong modulation and resonance at around 40 Hz, also known as the gamma frequency (Fries et al., 2007; Cardin et al., 2009; Lisman and Jensen, 2013). If responses at 40 Hz in one area are to be intelligible in another cortical or hippocampal area, then it is necessary for these circuits to be able to follow inputs at these frequencies. Here we showed that circuit-level FFEI inputs can in principle allow such rapid communication.

### Predictions

The FFEI models make a simple prediction: the absence of inhibition at cells receiving FFEI inputs should limit the ability of these cells to follow inputs at high temporal frequencies. An optogenetics approach could be used to control the activation of retinal ganglion cells (RGC) with high temporal precision, and the activity of LGN relay cells in response to RGC action potentials could be measured. A GABAzine infusion to block inhibition in the LGN should limit the ability of LGN relay cells to follow RGC inputs at high temporal frequencies. A similar experiment could be performed at thalamocortical slices in somatosensory cortex (Agmon and Connors, 1991) in order to test the high-convergence regime. While we have noted that there is a high degree of synaptic depression in thalamocortical synapses (Chung et al., 2002; Higley and Contreras, 2006), the data from an experiment should exhibit a strong shift, as in Figure 9, from the FFEI dynamics towards E-only dynamics.

## Conclusion

Feed-forward pairing of excitatory and inhibitory connections provides several functions for neural circuits. It helps to linearize computations that are impacted by rectification in retina (Werblin, 2010), LGN, and cortex (Hirsch et al., 2015). It sharpens stimulus tuning and spike timing (Pouille and Scanziani, 2001; Wehr and Zador, 2003; Blitz and Regehr, 2005; Mittmann et al., 2005; Higley and Contreras, 2006; Cardin, 2018), and allows neurons to respond to weak inputs without saturating for strong inputs (Pouille et al., 2009). In hippocampus, the delay between excitation and inhibition shortens with stronger CA3 inputs, producing smaller changes in CA1 membrane potential (Bhatia et al., 2019). Therefore feedforward excitation and inhibition allows neurons to accommodate a dynamic range of inputs before firing, providing a mechanism of gain control to these circuits (Pouille et al., 2009; Bhatia et al., 2019). Here we add that it provides a dual signal – the value of its input on the one hand and a clock-like trigger on the other – that allows high frequency feed-forward computation that is synchronized across brain areas.

## Impact statement

An archetypical feedforward projection pattern – excitation followed by a delayed inhibitory copy – can play the role of a clock-like trigger, allowing the organized processing of high frequency information within neural circuits.

## Acknowledgements

This work was funded by NIH EY022122 (SDV) and NIH DA033463. We thank Andrew Lipnick, whose undergraduate thesis on velocity tuning led us to consider feedforward transmission, Paul Miller for useful suggestions, and members of the Van Hooser lab for comments.

## Contributions

AC performed modeling, AC and SDV designed the models, AC and SDV wrote the paper.

---

**Supplementary Figure 1-1.**
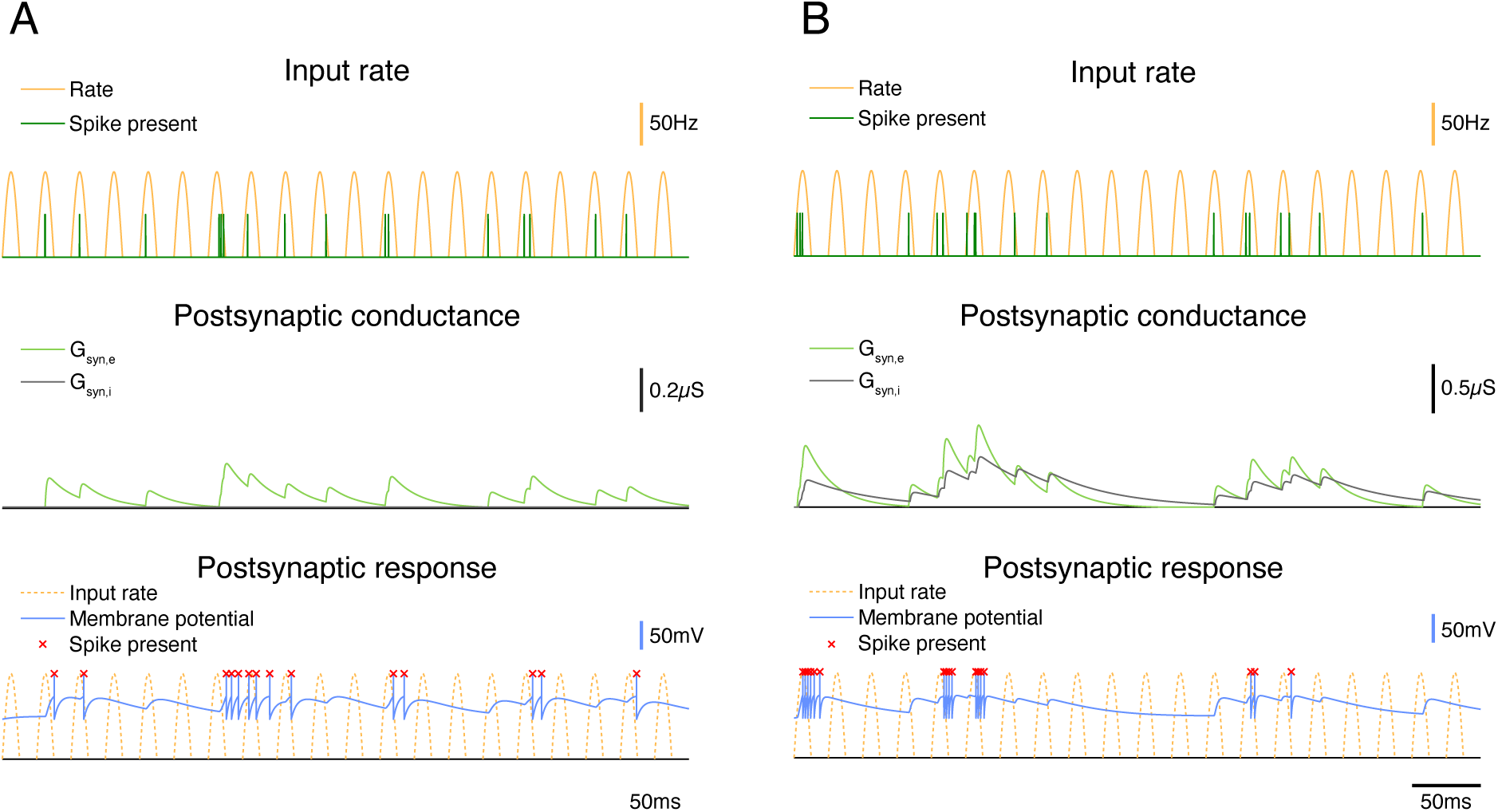
High frequency transmission differs between excitation-only circuits and circuits exhibiting paired feedforward excitation and delayed inhibition with long time constants. **(A)**: 0.5s simulation of a feedforward E circuit receiving a single rectified sinusoidal Poisson input (yellow) with a peak rate (PR) of 100 Hz and an input modulation frequency (F) of 40 Hz, as in Fig. 1A. **(B)**: Same, but for a paired feedforward E/I model circuit with τ_fall,i_ = 50 ms instead of the baseline 20 ms shown in Figure 1A. Spiking activity in the postsynaptic neuron with τ_fall,i_ = 50 ms is more prolonged than in the 20ms case in Figure 1A, but the firing is still truncated by the inhibition, as compared to the no-inhibition case in panel A.

**Supplementary Figure 6-1.**
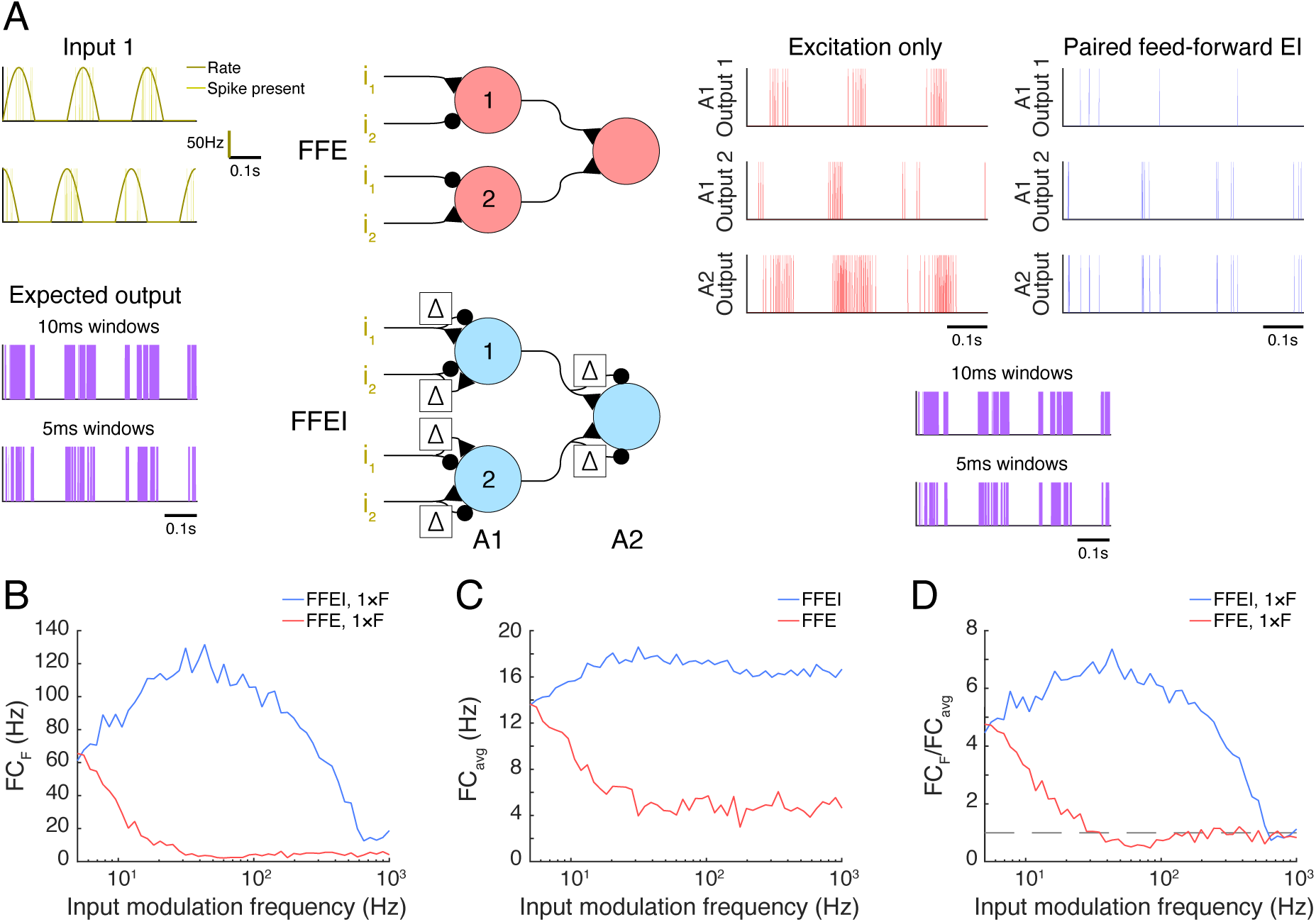
Paired feedforward E/I in an XOR circuit, with π/2 phase-shifted inputs. **(A)** A simulation of each XOR model circuit described in Figure 6A is shown with inputs modeled as in Figure 6A, but with a π/2 phase shift instead of a π phase shift, which should result in output at the stimulus frequency F (instead of 2 x F when the phase shift is π). **(B)** For a range of input modulation frequencies F, the Fourier coefficient of the A2 output at F (FC_F_) was averaged over 10 trials for the two XOR circuit models shown in **(A)**. **(C)** For a range of input modulation frequencies F, the mean power of the A2 output over all frequencies (FC_avg_). **(D)** For a range of input modulation frequencies F, the normalized Fourier coefficient of the A2 output is shown. Dashed line indicates FC_F_/FC_avg_ = 1.

**Supplementary Figure 8-1.**
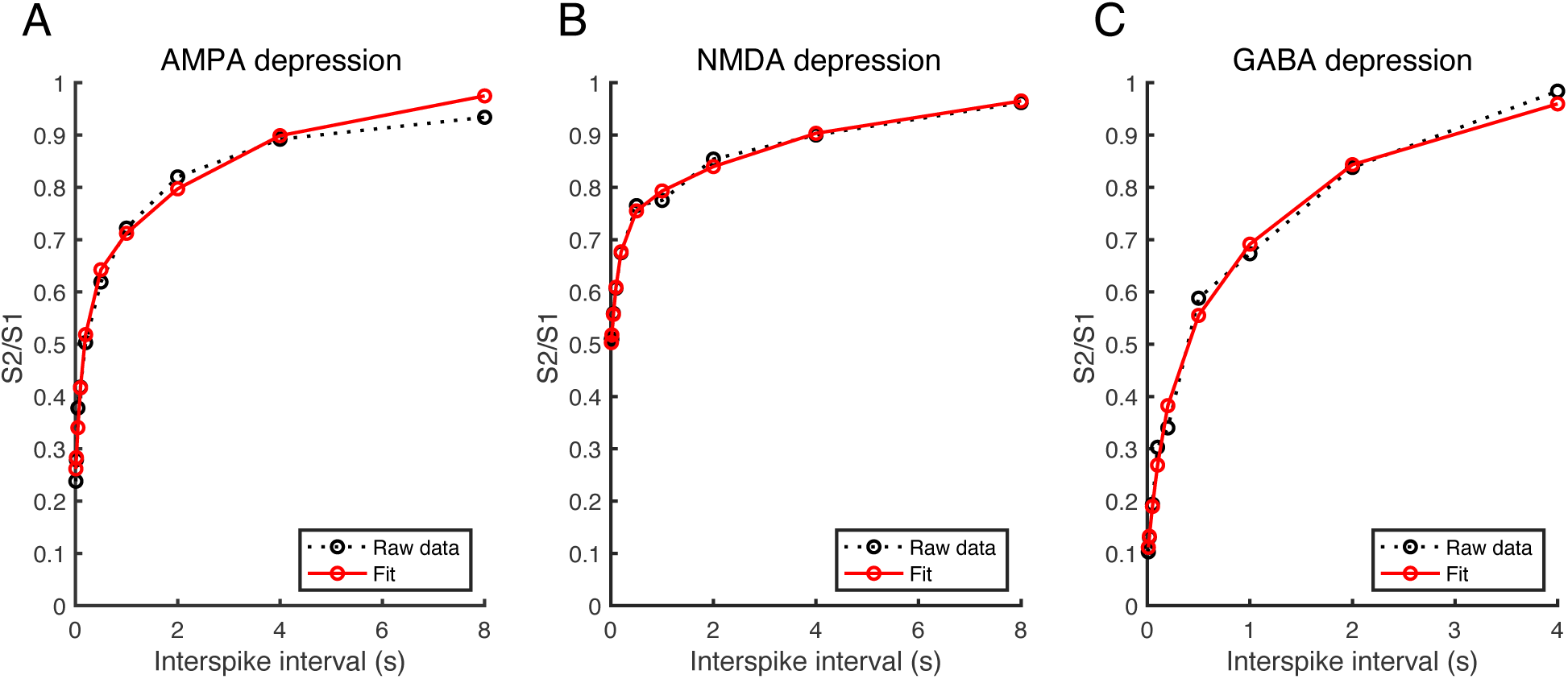
Fitting depression at the retinogeniculate synapse. **(A-C)** The paired-pulse ratio for the AMPAR **(A)**, NMDAR **(B)**, and locked GABAR **(C)** components of the synaptic current was plotted over a range of interspike intervals. For each channel, raw data (black) obtained from voltage clamp experiments (Chen et al., 2002; Blitz and Regehr, 2005) were fitted to a depression model (see Methods section). Paired-pulse experiments were then simulated using the fitted depression model (red).

